# Inhibition of Glycogen Metabolism Induces Reactive Oxygen Species-Dependent Apoptosis in Anaplastic Thyroid Cancer

**DOI:** 10.1101/2022.02.16.480663

**Authors:** Cole D. Davidson, Jennifer A. Tomczak, Eyal Amiel, Frances E. Carr

## Abstract

Anaplastic thyroid cancer (ATC) is one of the most lethal solid tumors, yet there are no effective, long-lasting treatments for ATC patients. Most tumors, including tumors of the endocrine system, exhibit an increased consumption of glucose to fuel cancer progression, and some cancers meet this high glucose requirement by metabolizing glycogen. Our goal was to determine if ATC cells metabolize glycogen and if this could be exploited for treatment. We detected glycogen synthase and glycogen phosphorylase (PYG) isoforms in normal thyroid and thyroid cancer cell lines and patient-derived biopsy samples. Inhibition of PYG using CP-91,149 induced apoptosis in ATC cells but not normal thyroid cells. CP-91,149 decreased NADPH levels and induced reactive oxygen species accumulation. CP-91,149 severely blunted ATC tumor growth *in vivo*. Our work establishes glycogen metabolism as a novel metabolic process in thyroid cells that presents a unique, oncogenic target that could offer an improved clinical outcome.

**Significance:** Glycogen metabolism plays an important role in combating reactive oxygen species and apoptosis in anaplastic thyroid cancer. Glycogen phosphorylase was inhibited with small molecule inhibitors that limited cell proliferation *in vitro* and blunted tumor growth in a nude mouse xenograft. This study demonstrates that glycogen metabolism is a viable target in one of the most lethal solid tumors.

## Introduction

Thyroid cancer is the most common malignancy of the endocrine system, and the incidence has greatly increased over the past forty years^1^. Well-differentiated thyroid cancers such as papillary (PTC) and low-grade follicular (FTC) thyroid cancers can usually be treated with surgery, radiation, or radioactive iodide (RAI) therapy. However, poorly differentiated (PDTC) and anaplastic thyroid cancers (ATC) are highly aggressive, metastatic, stemlike, and develop resistance to commonly used therapies^2^. Sorafenib is commonly used in ATC patients to target BRAFV600E, a gain of function mutation found in nearly 50% of ATC tumors^3^. Unfortunately, escape mechanisms arise that render sorafenib ineffective just months after application^4-7^. Therefore, there is an urgent need to address the abysmal clinical outcome for one of the most lethal tumors.

Historically, targeted therapies in poorly differentiated thyroid cancer and ATC have inhibited cell signaling kinases with varying levels of success. However, few strategies have been rigorously tested to target metabolic alterations in thyroid cancer^8^. Many cancer cells preferentially utilize glycolysis over oxidative phosphorylation even in the presence of oxygen, a phenomenon known as the Warburg effect^9^. The high rate of glycolysis in cancer cells may be to generate lactate to acidify the tumor microenvironment, favoring angiogenesis and immunosuppression, while generating nucleotides and lipids for cell division. Hexokinase has been identified as an attractive drug target to limit the first step in glycolysis. The hexokinase inhibitor 2-deoxyglucose (2-DG) reduced lactate production in an ATC xenograft but failed to significantly reduce tumor size^10^. Additionally, 2-DG treatment levies concern for long-term clinical application due to its broad-spectrum toxicity, particularly in the brain, heart, and stomach^11^. The pan-metabolic inhibitor metformin has been shown to limit ATC proliferation *in vitro* and is in a phase II clinical trial for differentiated thyroid cancer in combination with RAI^8,12,13^. However, metformin has many metabolic targets, and the therapeutic efficacy in cancer patients remains controversial^14,15^. Ideally, a metabolic inhibitor would exhibit low toxicity by inhibiting a metabolic pathway that is active only in a few cell types in normal tissue, in addition to the cancer cells being targeted.

In addition to importing glucose from the bloodstream, many tumors have been shown to fuel cell metabolism from storing and breaking down the glucose polymer, glycogen^16-19^. Glucose can be stored as a quick-access carbon cache via glycogen synthase 1 (GYS1) and rapidly broken back down to glucose via glycogen phosphorylase liver (PYGL) or brain (PYGB) isoforms. Glycogen deposits have been clinically observed in diverse types of cancer and have been shown to play a role in tumor progression^19-22^. Furthermore, inhibiting glycogen breakdown via RNAi has shown remarkable success in a glioblastoma xenograft model^19^. While glycogen represents a promising metabolic target in cancer, glycogen stores have only been reported in normal canine and bovine thyroids and the very rare clear cell thyroid carcinoma^23-25^. We recently hypothesized that thyroid cancer may utilize glycogen metabolism for tumor progression based on our own RNA-sequencing data and publicly available databases^26^. Therefore, our goal was to investigate the presence and role of glycogen in human thyroid tissues and cell lines and to determine the therapeutic potential of inhibiting glycogen metabolism.

## Results

### Glycogen Phosphorylase Brain Isoform Overexpression Drives Glycogen Breakdown in Thyroid Adenoma and Thyroid Cancers

We first profiled biopsy samples from normal thyroid and thyroid adenoma, PTC, FTC, and ATC to measure GYS1, PYGL, and PYGB expression. GYS1 expression was remarkably similar across the spectrum of thyroid cancers. GYS1 was slightly diminished in PTC and ATC compared to normal (Fig. 1A and 1D). PYGL expression did not change across the spectrum of thyroid differentiation (Fig. 1B and 1E). Surprisingly, PYGB was virtually undetectable in normal thyroid tissue, and was significantly upregulated in thyroid adenoma and all three forms of thyroid cancer (Fig. 1C and 1F). Each type of thyroid tissue likely had the potential to metabolize glycogen since all thyroid samples expressed GYS1 and at least one isoform of glycogen phosphorylase. We measured glycogen content using periodic acid–Schiff staining on the biopsy cores. As expected, glycogen content was inversely related to the presence of PYGB; normal thyroid tissue contained the most glycogen, followed by adenoma, PTC, FTC, and ATC (Fig. 1G-H), and a logarithmic relationship between glycogen staining and the sum of PYGL and PYGB expression was observed (Fig. 1I). Collectively, these results demonstrate that thyroid tissue, normal and malignant, across the spectrum of differentiation express enzymes necessary to store and breakdown glycogen.

**Figure 1.**
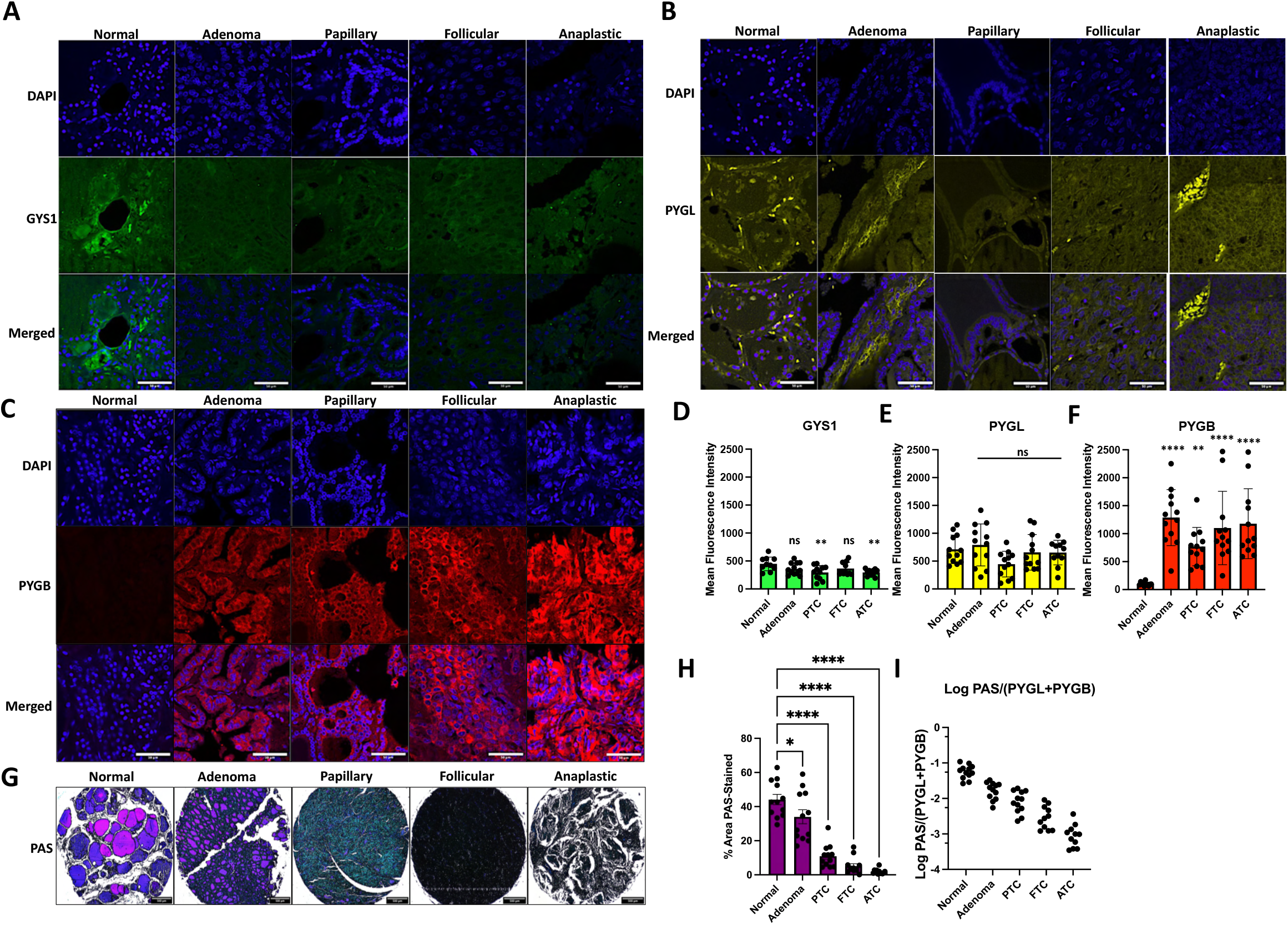
Glycogen metabolism enzymes are expressed in normal thyroid and thyroid cancer tissue from patient samples. **A-C**. Patient thyroid tissue microarrays were probed with antibodies for GYS1 (A), PYGL (B), and PYGB (C) and were stained with DAPI. Magnification = 20X, scale bar = 50 µm. **D-F**. Mean fluorescence intensity was calculated from each tissue core. **G**. Patient thyroid tissue microarrays were stained for glycogen content with PAS stain. Magnification = 4X, scalebar = 500 µm **H**. PAS stain was quantified as stain intensity per relative core area. **I**. PAS staining intensity was plotted as a logarithmic function with glycogen phosphorylase expression. N = 11-12 for each type of thyroid classification.

### Representative Cell Lines of Normal Thyroid and Thyroid Cancer Cells Metabolize Glycogen through Expression of Glycogen Synthase and Glycogen Phosphorylase Isozymes

We next tested thyroid cell lines to directly investigate the presence and role of glycogen in normal thyroid and thyroid cancer. We used cell lines representing normal thyroid, PTC, FTC, and ATC from diverse genetic backgrounds (Supplementary Table S1)^27^. On the RNA level, *GYS1* expression was comparable across all cell lines except for the OCUT-2 cell line that expressed nearly four times more *GYS1* transcript than the normal thyroid cells, Nthy-ori-3-1 (Fig. 2A). This was not unexpected, as GYS1 overexpression has been reported in many cancer cell models^28,29^. Additionally, *PYGL* expression was comparable in the different types of thyroid cancer cells except the FTC-133 and 8505C cells expressed significantly higher levels (Fig. 2B). In agreement with the biopsy results, *PYGB* expression correlated with thyroid dedifferentiation; we observed the highest *PYGB* expression in the ATC cell lines OCUT-2 and 8505C (Fig. 2C). Immunoblot analysis of glycogen metabolism enzymes (Fig. 2D) revealed differences in levels of phosphorylated (deactivated) GYS1. Interestingly, normal, PTC, and an ATC cell line displayed heavily phosphorylated GYS1, whereas both FTC cell lines and the ATC cell line, OCUT-2, displayed decreased GYS1 phosphorylation. The complicated profile is likely reflective of the genetic background of each cell line, as FTC-133, CUTC61, and OCUT-2 all have mutations in the PI3K-Akt-GSK3β pathway, which results in reduced GYS1 phosphorylation^30^. Overall, the thyroid cell line profiles are consistent with the patient tissue staining; PYGB is overexpressed in thyroid cancer. The dynamic expression and activation of glycogen enzymes may impact the differential levels of glycogen in the cell lines, as TPC-1, FTC-133, and OCUT-2 had the highest levels of glycogen (Fig 2E). We validated our detection of glycogen in the Nthy-ori-3-1 and 8505C cells using tannic acid staining and transmission electron microscopy (TEM). We observed large (∼45 nm), organized glycogen particles in the normal thyroid cells (Fig. 2F) compared to the smaller (∼25 nm), dispersed glycogen particles in the 8505C cells (Fig. 2H). Glycogen became nearly undetectable following overnight glucose starvation in both cell lines (Fig. 2G and 2I), insuring staining specificity.

**Figure 2.**
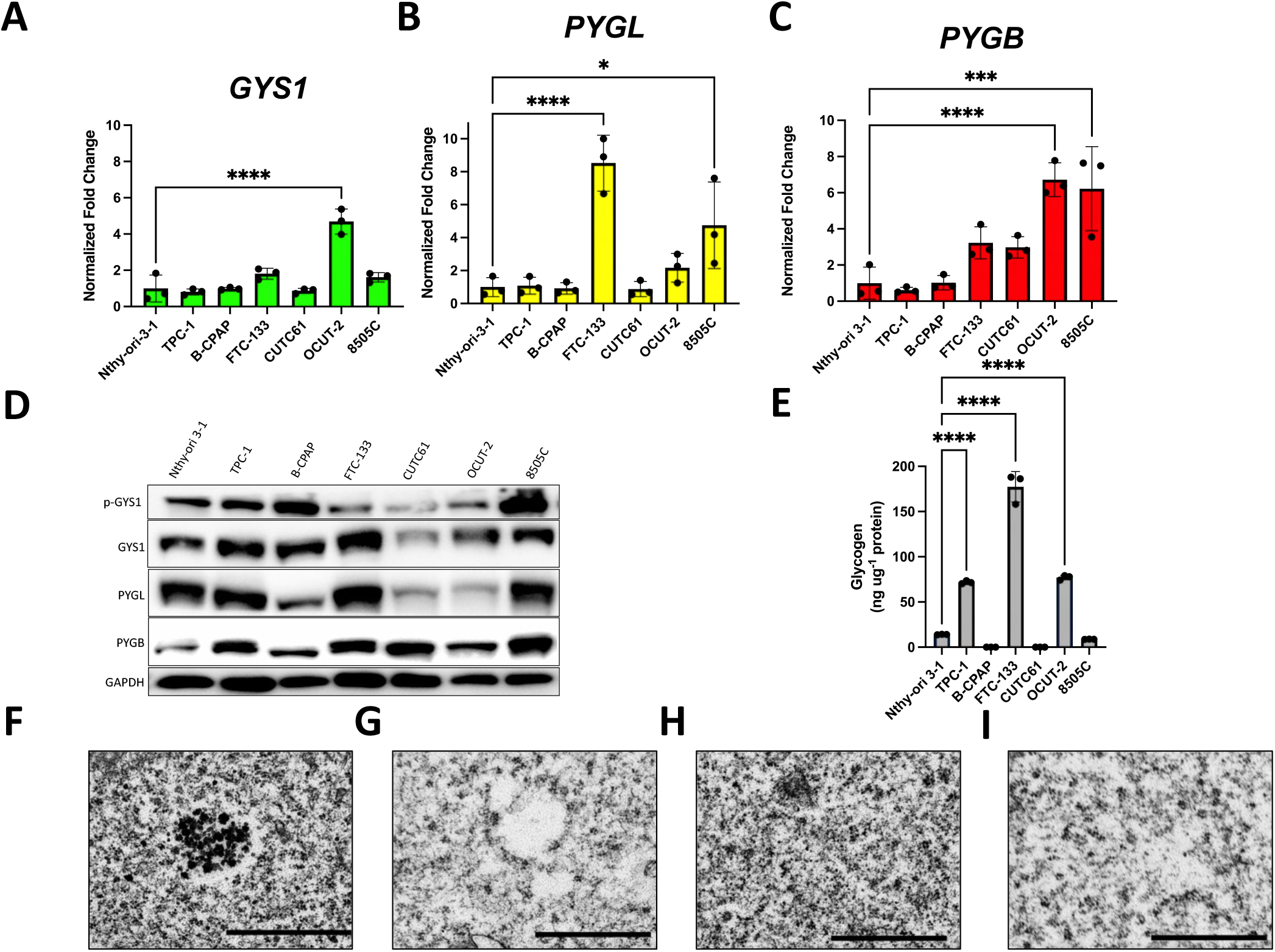
Representative normal thyroid and thyroid cancer cell lines express the glycogen machinery necessary to synthesize and store glycogen granules. **A-C**. Normal thyroid and thyroid cancer cell lines express *GYS1* (A), *PYGL* (B), and *PYGB* (C) as determined by RT-qPCR. **D**. Thyroid cell lines were profiled for protein expression of phosphorylated GYS1, total GYS1, PYGL, and PYGB. **E**. Glycogen content was measured in each thyroid cell line via colorimetry. **F-I**. Transmission electron microscopy revealed glycogen deposits in Nthy-ori-3-1 (F) and 8505C cells (H) that were reduced via overnight glucose starvation (G and I). Magnification = 5000X, scalebar = 1 µm.

### Glycogen Phosphorylase Inhibition Increases Glycogen Content to Limit Cell Viability and Proliferation and Induce Apoptosis in ATC Cells

Following the observations that thyroid cancer cells store glycogen and overexpress PYGB, we reasoned that glycogen phosphorylase could be a promising drug target, particularly in ATC cells for which there is an urgent need for effective, long-term therapeutics. We focused on the 8505C cells, which are well representative of ATC cells, having no mutations in *PIK3CA* and low levels of glycogen (Fig. 1G-H)^31^. CP-91,149 (CP) was originally designed to treat diabetes and is a pan glycogen phosphorylase inhibitor by stabilizing the enzyme homodimer in an inactive form^32,33^. CP significantly increased glycogen content in normal and ATC cells as shown through colorimetry and TEM (Fig. 3A-E). CP also modestly but significantly reduced total cell viability in 48 hours at 50 µM in the ATC cells but not the normal thyroid cells (Fig. 3F, Supplementary Fig. S1A-D). Inhibiting PYG with CP has been shown to induce intrinsic apoptosis in hepatocellular carcinoma^21^, so we were prompted to investigate the mechanism of cell death in thyroid cells. CP caused significant PARP, caspase 9, and caspase 7 cleavage in 8505C cells in only six hours (Fig. 3G and 3I-K). Additionally, incubation with CP resulted in reduced expression of the proliferation marker Ki-67 (Fig. 3G and 3L). Conversely, CP did not significantly induce apoptosis in Nthy-ori-3-1 cells as observed by lack of PARP and caspase cleavage (Fig. 3H and 3M-O). Proliferation was also not affected in Nthy-ori-3-1 cells as evident from Ki-67 levels (Fig. 3H and 3P), suggesting that ATC cells are more sensitive to glycogen inhibition than normal thyroid cells. Furthermore, 24 hours of CP treatment inhibited the ability of 8505C cells to attach to fibronectin-coated plates (Supplementary Fig. S1E-F). To validate the specificity of CP-91,149, we employed siRNA targeting *PYGB* in 8505C cells (Supplementary Fig. S2A-C). We observed a 50% increase in glycogen following *PYGB* knockdown, as well as a significant decrease in cell viability (Fig. 3Q-R). We also assessed CP efficacy on ATC stem cells, which is an important consideration as many inhibitors often fail to kill the pluripotent stem cell population^34,35^. Excitingly, 8505C stem cells exhibited a concentration-dependent response to CP (Fig. 3T-U). This observation may be reflective of stem cells’ higher energy requirement and dependence on glycogen as seen in undifferentiated human embryonic stem cells and mesenchymal stem cells^36,37^. In addition to PYG inhibition with CP-91,149, ATC cells were sensitive to CP316819. CP316819, like CP-91,149, is an indole-ring containing small molecule inhibitor of PYG that enhanced glycogen storage in rat brains^38^. CP316819 limited cell viability and increased the glycogen content of 8505C cells (Supplementary Fig. S3A-D).

**Figure 3.**
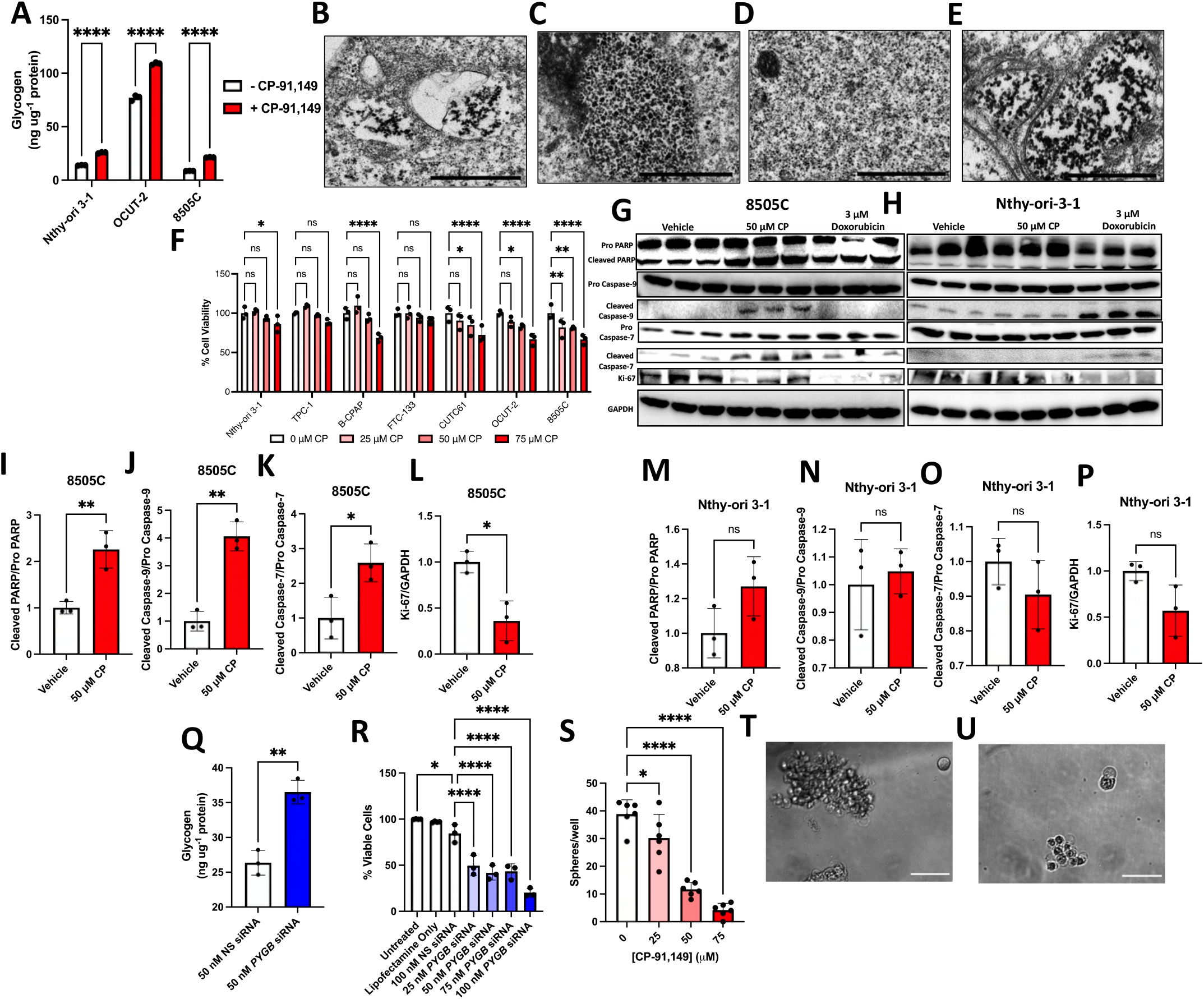
Glycogen phosphorylase inhibition induces glycogen buildup and apoptosis in anaplastic thyroid cancer cells but not normal-like thyroid cells. **A-E**. CP-91,149 (50 µM, 24-hour incubation) increased glycogen content in Nthy-ori-3-1 and ATC cells as shown by colorimetry (A) and TEM (B-E). Magnification = 5000X, scalebar = 1 µm **F**. Thyroid cells exhibit a concentration-dependent decrease in cell viability to CP-91,149 (48 hour incubation). **G-H**. CP-911,49 induces apoptosis and inhibits proliferation in 8505C (G) but not Nthy-ori-3-1 cells (H) as shown via cleaved PARP, cleaved caspase 9 and 7, and Ki-67 levels. **I-P**. Quantification of immunoblots for apoptosis markers and Ki-67. **Q**. Knockdown of *PYGB* is sufficient to induce glycogen buildup in 8505C cells. **R**. 8505C cell viability is reduced following three days of *PYGB* knockdown. **S-U**. CP-91,149 inhibits 8505C thyrosphere growth at day 3 (U) compared to day 1 (T). Magnification = 40X, scalebar = 100 µm.

### ATC Cells are Sensitive to Glycogen Synthase Inhibition

Following the recent success of using naturally occurring inhibitors of GYS1 for treating adult polyglucosan body disease^39^, we treated 8505C and OCUT-2 cells with the flavoring agent guaiacol to decrease ATC cell viability and glycogen levels (Supplementary Figure S3E-G). We also used the recently characterized molecule yGsy2p-IN-1, the first specific GYS1 inhibitor^40^, to limit ATC viability and decrease glycogen content (Supplementary Figure S3H-I). These data may be the first report of using guaiacol or yGsy2p-IN-1 in cancer cell models to limit cell viability by blocking glycogen utilization, highlighting the potential of targeting glycogen metabolism in aggressive cancers.

### CP-91,149 Triggers Glucose Flux in ATC Cells to Fuel Glycolysis but Inhibits NADPH Production, Increases Levels of Reactive Oxygen Species, and Limits Oxidative Phosphorylation

Glycogen provides efficient access to glucose for the cancer cell to use in diverse cellular processes. We first confirmed that CP resulted in a buildup of the glycogen monomer, glucose-1-phosphate (G1P) and a decrease in the glycolytic intermediate, glucose-6-phosphate (G6P) in 8505C cells (Fig. 4A-B). A characteristic of high glycolytic activity is lactate production, which acidifies the extracellular environment. While 2-DG significantly inhibited the extracellular acidification rate (ECAR), we were surprised to find that CP modestly but significantly enhanced ECAR in 8505C cells (Fig. 4C-D). These results were confirmed with an extracellular lactate assay (Fig. 4E). We reasoned that this paradox could be explained by a “glucose stress response” in the cell, in which glucose intake was increased to compensate for glycogen catabolism inhibition. We chose to measure expression of the glucose transporters most implicated in ATC; GLUT1, GLUT3, and GLUT4 (Supplementary Figure S4A-C)^41-43^. There was an increase in GLUT1 and GLUT3 (*SLC2A1* and *SLC2A3*) expression following CP treatment, while the glucose-independent GLUT4 (*SLC2A4)* expression did not change (Fig. 4F-H). There was a significant reduction in extracellular glucose following CP treatment as well (Fig. 4I). Unexpectedly, the ECAR readout with 2-DG treatment was equivalent regardless of CP pretreatment (Fig. 4C-D). We previously showed that free glucose and glycogen stores made distinct contributions to ECAR in murine dendritic cells, which suggests that glycogen may not contribute to the glycolytic glucose pool in ATC cells^44^. Therefore, we investigated function of the pentose phosphate pathway (PPP) following glycogen phosphorylase inhibition. The PPP siphons G6P from glycolysis or glycogenolysis to reduce NAD+ to NADPH via glucose-6-phosphate dehydrogenase (G6PDH). NADPH is a crucial redox metabolite in cancer biology for combating reactive oxygen species (ROS) produced in metabolism and cell signaling^45^. There was a drastic decrease in NADPH in cells treated with CP compared to control cells (Fig. 4J), which has important implications for the redox potential. Since proteins and nucleotides oxidized by ROS require proteins such as glutathione for repair, we measured reduced and total glutathione following CP treatment and observed a concomitant loss in reduced glutathione (GSH) (Fig. 4K). We measured a three-fold increase in ROS in ATC cells following CP treatment, which was ablated with exogenous NADPH or the glutathione precursor N-acetylcysteine (NAC) (Fig. 4L). To confirm the mechanism of CP-mediated loss in cell viability, we partially rescued CP-treated 8505C cells with NADPH, NAC, and a manganese superoxide mutase mimetic (Mn-TMP) (Fig. 4M). Since high levels of ROS can impair mitochondrial activity and induce apoptosis^46-48^, we measured the oxygen consumption rate (OCR) following CP treatment. As expected, CP significantly limited the OCR in ATC cells (Fig. 4O). Intriguingly, vehicle-treated ATC cells could compensate for inhibition of glycolysis with 2-DG by increasing OCR, but the CP-treated cells were unable to increase their oxidative phosphorylation (Fig. 4P).

**Figure 4.**
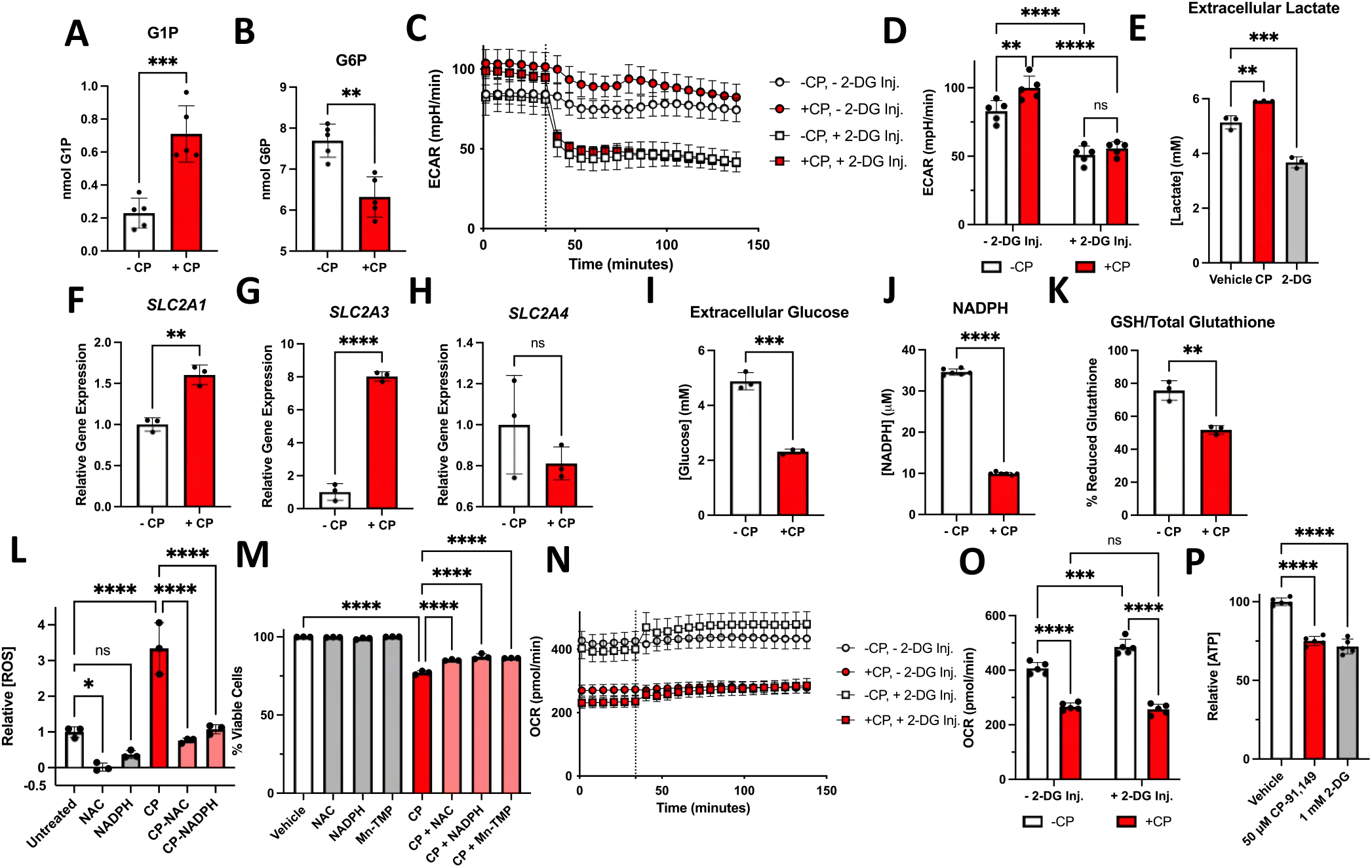
Glycogen phosphorylase inhibition induces disruption in glycolysis, the pentose phosphate pathway, reactive oxygen species levels, and oxidative phosphorylation. **A-B**. CP-91,149 causes a buildup in glucose-1-phosphate (A) and a reduction in glucose-6-phosphate (B). **C-E**. CP-91,149 enhances extracellular acidification rate (C-D) and lactate production (E). **F-H**. CP-91,149 treatment increases the expression of *SLC2A1*, (GLUT1) (F), *SLC2A3*, (GLUT3) (G), and *SLC2A4*, (GLUT4) (H). **I**. CP-91,149 causes an increase in glucose uptake from the cell culture media. **J-K**. CP-91,149 decreases the available supply of NADPH (J) and decreases the relative amount of reduced glutathione (GSH) (K). **L**. CP-91,149 enhances the relative levels of reactive oxygen species which can be attenuated with exogenous N-acetyl cysteine or NADPH. **M**. Exogenous N-acetyl cysteine, NADPH, and MN-TMP can rescue ATC cells from CP-mediated loss in cell viability. **N-O**. CP-91,149 decreases the basal oxygen consumption rate in ATC cells. **P**. CP-91,149 treatment decreases the levels of ATP in ATC cells. 8505C cells were incubated with 50 µM CP-91,149 for 24 hours prior to all experiments.

Expanding on the potential for separate glucose pools and functions, we modulated levels of glucose and pyruvate in combination with CP and 2-DG treatment. Since both CP and 2-DG limit the amount of G6P to enter glycolysis or the PPP, it was expected that depriving cells of glucose would decrease cell viability in both treatment groups (Supplementary Figure S5A). Excess (25 mM) glucose rescued cell viability after treatment with CP or 2-DG, suggesting that exogenous glucose can compensate loss of both glycolysis and glycogenolysis activity. However, ROS levels were higher in cells treated with CP compared to 2-DG, and exogenous glucose only rescued ROS levels induced by CP (Supplementary Figure S5B). Unlike CP, 2-DG only modestly increased ROS levels, and while excess glucose could rescue the viability of cells treated with 2-DG or CP, excess glucose reduced ROS levels only in CP-treated cells. Conversely, supplementing the PPP with exogenous NADPH reduced ROS in both treatment groups (Fig. 4L and Supplementary Fig. S5C) but only rescued cell viability in the CP-treated cells (Fig. 4M and Supplementary Fig. S5D). Taken together, these findings suggest that exogenous glucose restores functionality to glycolysis, not the PPP in 2-DG treated cells, and exogenous NADPH restores functionality to the PP in both treatment groups but is only sufficient to rescue viability in CP-treated cells. Pyruvate depletion only significantly impacted the 2-DG-treated cells, suggesting that the cells treated with 2-DG were more dependent on the high energy pyruvate, while the CP-treated cells were able to compensate for loss of pyruvate (Supplementary Figure S5A). Likewise, only the 2-DG-treated cells could be rescued with excess (2 mM) pyruvate, whereas the CP-treated cell viability level was nearly identical to the pyruvate-depleted group. These findings support a differential mechanism of CP on cell viability compared to 2-DG.

### CP-91,149 Exhibits Drug Synergy with Inhibitors of MAPK Signaling and the Pentose Phosphate Pathway

Following the promising *in vitro* applications of CP in ATC cells, we evaluated the potential for CP to improve the efficacy of other inhibitors by performing cell viability assays to calculate coefficient of drug interaction (CDI) scores to assess drug synergy, additivity, and antagonism. Excitingly, CP displayed a high degree of drug synergy with the multi kinase inhibitor sorafenib (Fig 5A-B). Since sorafenib resistance often develops in ATC tumors, combining the kinase inhibitor with a metabolic inhibitor such as CP could yield more promising results, possibly at lower doses. Interestingly, CP displayed drug additivity rather than synergy with the multi kinase inhibitor lenvatinib (Fig. 5C-D), which is approved for aggressive differentiated thyroid cancer^49^ and the PI3K inhibitor buparlisib (Fig. 5E-F), which is in phase II trials for dedifferentiated thyroid cancer^50^. While MAPK signaling does not directly affect glycogen metabolism, both buparlisib and lenvatinib directly target upstream regulators of GYS1, resulting in slight drug profile overlap with CP. Furthermore, CP did not achieve synergy with the cell cycle inhibitor palbociclib or the topoisomerase inhibitor doxorubicin (Fig. 5G-J). We also performed synergy studies with other metabolic inhibitors to further elucidate the mechanism of cell death following CP treatment. Combination of CP with 2-DG resulted in CDI values close to 1.0 (Fig. 5K-L), which suggests drug additivity and redundant inhibition, expected from limiting G6P production from both agents. An alternative hexokinase inhibitor, 3-bromopyruvic acid (3-BP), also failed to achieve synergy with CP (Fig. 5M-N). We then inhibited the PPP directly using the G6PDH inhibitor, 6-aminonicotinamide (6-AN). Unlike glycolysis inhibitors, CP and 6-AN achieved remarkable drug synergy (Fig. 5O-P), potentially by inhibiting G6PDH activity and substrate availability in concert. Substituting CP for 6-AN support these findings; 6-AN and 2-DG displayed drug antagonism while 6-AN and sorafenib exhibited drug synergy (Supplementary Figure S6A-B).

**Figure 5.**
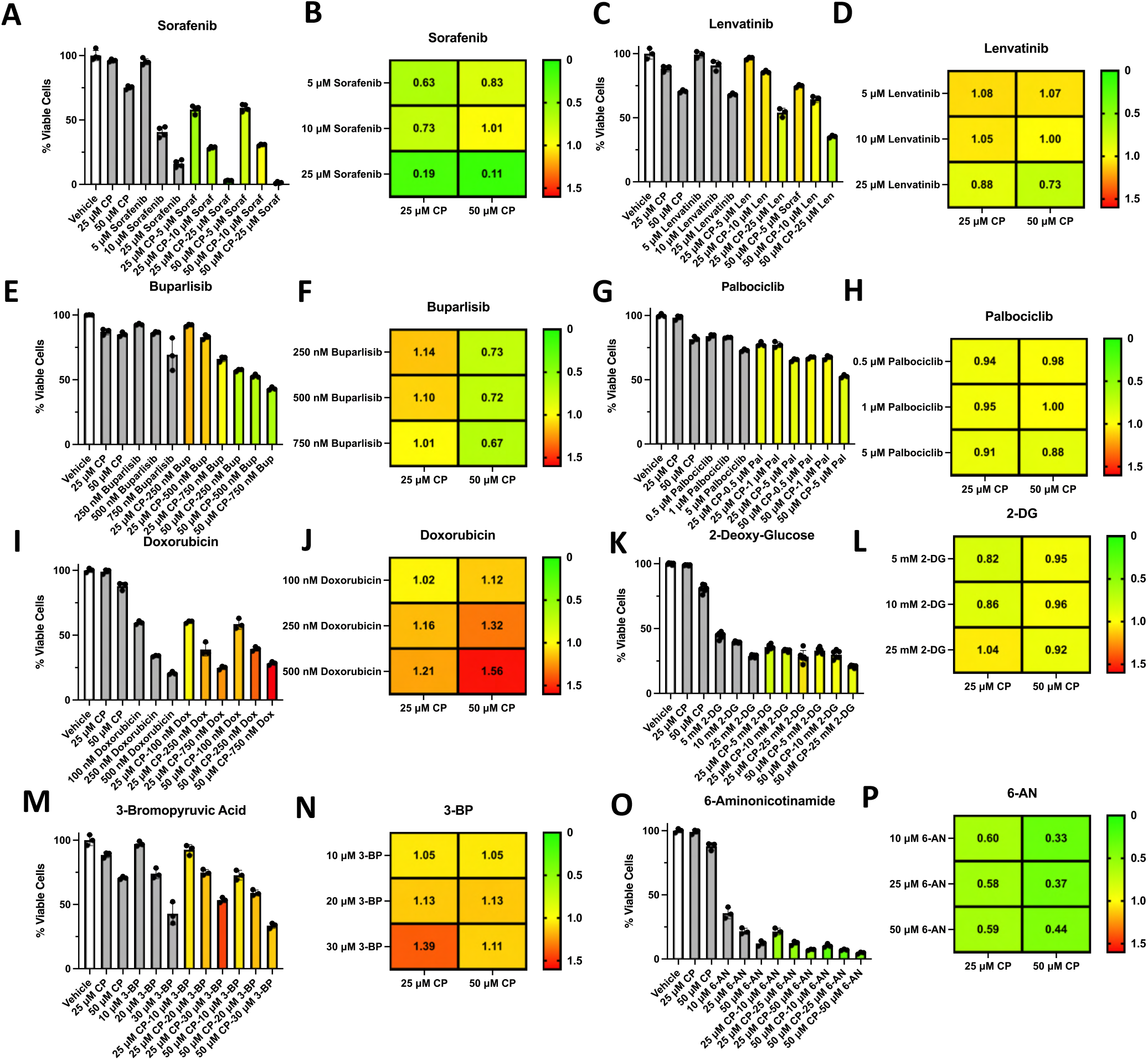
CP-91,149 displays differential drug additivity and synergy profiles with a diverse array of therapeutics. **A-P**. ATC cell viability was determined with CP-91,149 alone and in combination with sorafenib (A-B), lenvatinib (C-D), buparlisib (E-F), palbociclib (G-H), doxorubicin (I-J), 2-deoxy-glucose (K-L), 3-bromopyruvic acid (M-N), and 6-aminonicotinamide (O-P) to calculate coefficient of drug interaction scores.

### CP-91,149 Induces Apoptosis *in vivo* to Restrict ATC Tumor Growth

In order to validate our cell-based findings *in vivo*, nude mice were injected subcutaneously with 8505C in each flank to monitor tumor growth. Once palpable tumors formed (∼100 mm^3^, 10 days post injection), mice were given intraperitoneal injections of vehicle, CP, sorafenib, or combination of CP and sorafenib 6-7 days a week. Tumor growth was notably stunted at approximately equal levels in the CP, sorafenib, and combination group (Fig. 6A-D). Although the potential for drug synergy was masked due to the equivalent efficacy with sorafenib, the results demonstrate that CP was equally as effective as sorafenib. Importantly, CP displayed low toxicity, as all mice steadily gained weight over the course of treatment, demonstrating the potential for clinical translation (Fig. 6E). Following completion of the xenograft, tumors were excised and prepared for immunofluorescence experiments for proliferation and apoptosis. Xenografts from mice treated with CP alone or in combination with sorafenib showed a reduction in Ki-67 expression (Fig. 6F-G), and CP profoundly induced apoptosis as evident by the amount of PARP cleavage (Fig. 6F and H). In agreement with the *in vitro* data, CP significantly increased the level of glycogen in the tumor tissue (Fig. 6F and I). Surprisingly, the combination treatment significantly decreased KI-67 expression but did not demonstrate a significant increase in PARP cleavage or glycogen staining (Fig. 6F-I). This could be reflective of late-stage apoptosis or even necrosis where cleaved PARP and glycogen are fully degraded by lysosomal hydrolases^51^. While further studies will be required to investigate specifically how environmental factors such as diet and exercise could impact a glycogen-targeted therapeutic regimen, these data underscore the strong potential for cancer strategies targeting glycogen phosphorylase.

**Figure 6.**
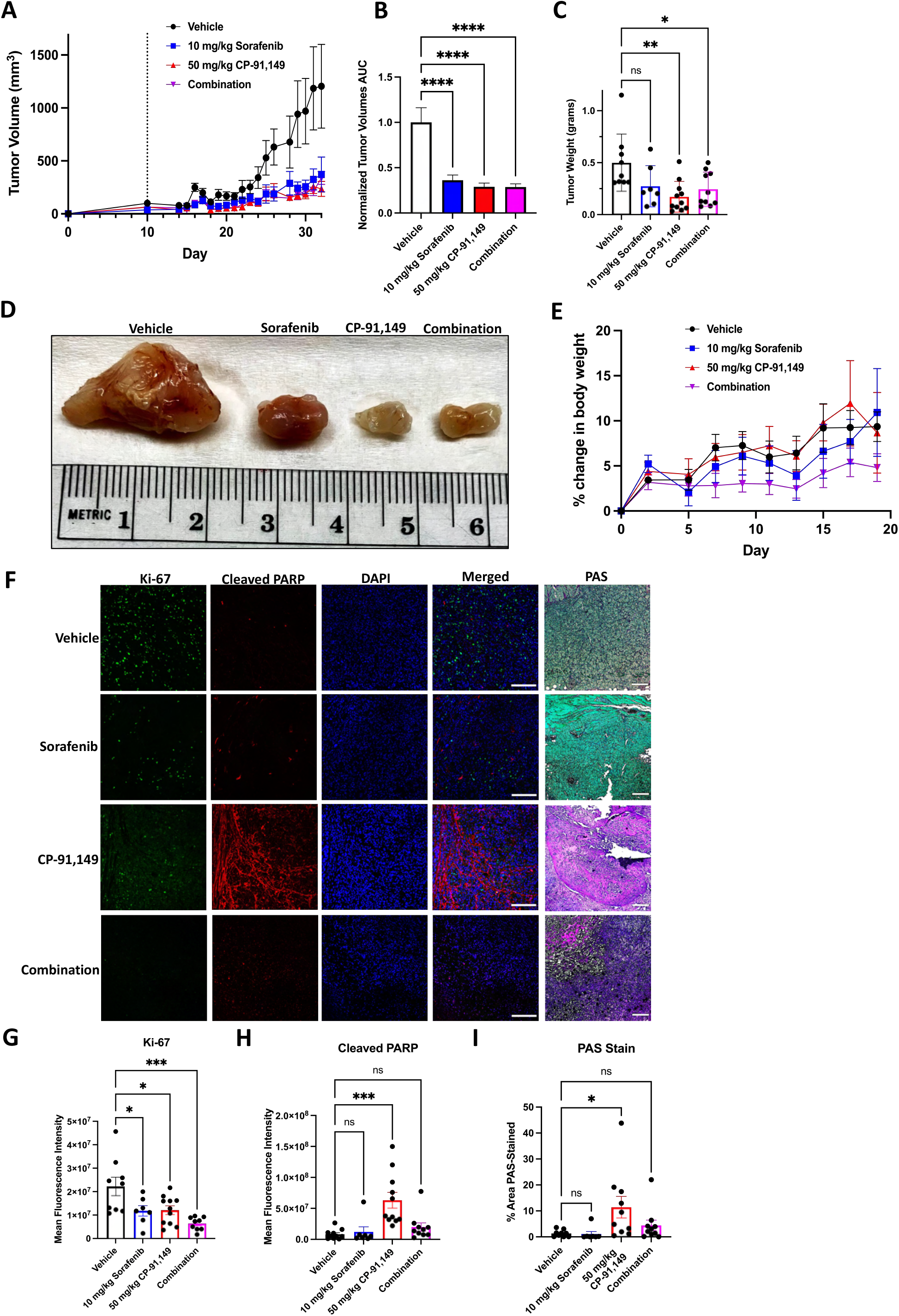
CP-91,149 significantly inhibits tumor growth and induces apoptosis in an ATC xenograft model. **A**. Nude mice were injected subcutaneously with 8505C cells on day 0, and tumors were allowed to grow for 9 days when mice were reassigned cages to achieve approximately equal weights in each treatment group. Mice were injected daily for 22 days beginning on day 10 (dashed line) with vehicle control, 10 mg/kg sorafenib, 50 mg/kg CP-91,149, or a combination of sorafenib and CP. **B**. Relative area under the curve for each treatment group. **C**. Mice were sacrificed on day 33 and tumors were excised and weighed. **D**. Representative image of an excised tumor from each treatment group. **E**. Mice were weighed three times a week following drug injections to monitor toxicity. **F**. Representative images of tumors stained for Ki-67, cleaved PARP, and glycogen. 20X, scalebar = 150 µM for IF. 4X, scalebar = 500 µM for PAS. **G**. Quantification of Ki-67 fluorescent intensity. **H**. Quantification of cleaved PARP fluorescent intensity. **I**. Quantification of percent of sample area stained with PAS.

## Discussion

Successful tumor progression relies on metabolic reprogramming for enhanced energy reserves and a larger supply of metabolic building blocks for daughter cells. Nearly all cancer cells exhibit the Warburg effect to meet these high metabolic needs^9^. Unfortunately, directly inhibiting glycolysis with small molecule inhibitors such as 2-DG or 3-BP have resulted in unacceptable side effects^11,52^. Therefore, it may be more advantageous to target an altered metabolic pathway in cancer cells that is not active in every cell, such as glycogen metabolism.

Glycogen metabolism is emerging as an important process in cancer biology, as it has been found to play oncogenic roles in diverse tumor types. There are conflicting reports in the literature on the role and quantity of tumor glycogen compared to healthy, normal tissue counterparts. For example, liver cells, which rely on glycogen for gluconeogenesis to supply the body with glucose, have significantly less glycogen than liver cancer cells^16^. On the other hand, cancer cells from tissues normally devoid of glycogen appear to upregulate glycogen stores^17^. Perhaps the role and abundance of glycogen in specific cancer types are linked to the level of dedifferentiation; since healthy liver cells contain the most glycogen of any human cell, liver cancer cells subvert glycogen functionality to instead supply only the cancer cell with glucose. Conversely, normal breast cells are low in glycogen while breast cancer cells may begin to synthesize glycogen for times of low nutrient viability and hypoxia^18,19^. We found that normal thyroid tissues are rich in glycogen (Fig. 1G), and glycogen stores decrease with each step of dedifferentiation towards ATC progression (Fig. 1H). This raises the provocative implication that glycogen may have a physiological role in the thyroid and an oncogenic role in thyroid cancer. The cytoplasm of normal thyrocytes is one of the most oxidative environments in the body due to the production of H_2_O_2_ to oxidize dietary iodide^53^. In agreement with our observations that inhibition of glycogenolysis results in ROS accumulation in ATC cells (Fig. 4L), the role of glycogen in normal thyroid cells could be to protect against ROS formed during thyroid hormone synthesis. Further studies will be required to determine the exact fate of glycogen-derived carbon in thyroid cancer compared to normally functioning thyrocytes.

Although glycogen anabolism and catabolism represent exciting drug targets in cancer biology, few studies have investigated this potential. CP-91,149 was originally developed to treat diabetes by binding to the indole-site of glycogen phosphorylase, blocking the homodimer interface^32^. CP-91,149 inhibits all PYG isoforms and crosses the blood brain barrier for potential application in treating brain cancers. However, PYG inhibitors could be better optimized to suit the metabolic profile of different tumors. For example, potential toxicity could be avoided by exclusively targeting PYGB over PYGL. Nevertheless, no side effects have been reported following CP-91,149 treatment in mice; 50-100 mg/kg of CP-91,149 decreased obese mouse blood glucose levels but not those of lean mice, and we report here that all mice steadily gained weight regardless of treatment (Fig. 6E).

Inhibiting glycogen phosphorylase has garnered exciting *in vitro* results in diverse cancer cell lines. Pancreatic adenocarcinoma cells were found to be sensitive to CP-320626, a PYG inhibitor similar in structure to CP-91,149^20^. CP-320626 inhibited cellular respiration, the PPP, anaplerosis, and synthesis of ribose and fatty acids, indicating a major redistribution of carbon due to glycogenolysis inhibition. Studies in cell lines from glioblastoma, breast cancer, and colon cancer revealed that GYS1 and PYGL expression were increased in response to hypoxia^19^. Furthermore, knocking down *PYGL* induced ROS accumulation and senescence in cancer cells. In hepatocellular carcinoma, CP-91,149 also increased ROS levels but triggered autophagic adaptations^21^. In agreement with our findings (Fig. 5A-B and 5K-N), CP-91,149 treatment in HepG2 cells displayed synergy with sorafenib but not 2-DG or 3-BP. Fewer studies have been conducted on inhibiting glycogen synthase in cancer, as most pharmacological strategies target the upstream regulator of GYS1, glycogen synthase kinase 3. Guaiacol, a naturally occurring flavoring agent, was recently identified as a direct inhibitor of GYS1 by competing with the substrate UDP-glucose for incorporation into the growing glycogen chain^39^. Guaiacol decreased glycogen stores in mouse embryonic fibroblasts by 50%, which is comparable to guaiacol treatment in ATC cells (Supplementary Fig. S3G). A recent synthetic GYS1 inhibitor, yGsy2p-IN-1, robustly inhibited purified GYS1 and decreased ^14^C-glucose incorporation in cell lysates^40^. We report here that yGsy2p-IN-1 successfully reduced glycogen levels and limited cell viability in ATC cells (Supplementary Fig. S3H-I), demonstrating the inhibitor’s antimetabolic effects *in vitro*. Further studies are needed to better elucidate optimal drug combinations with a glycogen metabolism modulator based on the genetic background and cell signaling landscape of individual tumors.

Although ATC is one of the most lethal solid tumors with no long-term therapeutic option, no study thus far has directly investigated glycogen in normal thyroid or thyroid cancer cells. We report that normal thyroid tissue as well as thyroid adenoma, PTC, FTC, and ATC express the genes and enzymes necessary to metabolize glycogen (Fig. 1-2). Interestingly, normal thyroid tissue expressed PYGL but not PYGB. While PYGL and PYGB exhibit comparable enzyme kinetics, they have different mechanisms of allosteric regulation. PYGL is more dependent on hormone signaling from insulin and glucagon, while PYGB activity is more impacted through intracellular nutrient availability, e.g., G6P and AMP/ATP ratio^54^. PYGB may confer an oncogenic advantage over PYGL; instead of responding to hormones to selflessly catabolize glycogen for the body, PYGB is expressed to directly serve the metabolic needs of the cancer cell. Since PYGB overexpression has been observed in several aggressive tumors such as colorectal carcinoma, non-small lung cancer, metastatic breast cancers, and now anaplastic thyroid cancer, PYGB expression and the concomitant shunt in glycogen catabolism may represent a crucial step in successful tumor progression^55-57^. The levels of glycogen in the thyroid tissue samples were depleted with PYGB expression, which can be inhibited with small molecule inhibitors to cause glycogen buildup (Fig. 3A-E). A lack of available glycogen causes a depletion of NADPH and GSH, a rise in ROS levels, impaired OCR, and apoptosis induction (Fig 3G, 4J-K, and 4N-O). Based on these results, we propose that glycogen-derived carbon preferentially fuels the PPP over glycolysis (Figure 7). Further studies are needed to investigate the exact mechanism of directing this differential carbon trafficking. One explanation may be that the colocalization of glycogen granules and enzymes known as the glycosome is localized to the endoplasmic reticulum. Glycogen has been shown to be associated with ER membranes, and cancer cells rely heavily on the PPP in the ER to combat the unfolded protein response^58-60^. Excitingly, CP-91,149 achieved profound drug synergy with sorafenib *in vitro* and significantly impaired tumor growth *in vivo* (Fig 5A and 6). Taken together, these results expose glycogen metabolism to be a novel metabolic vulnerability in ATC cells.

**Figure 7.**
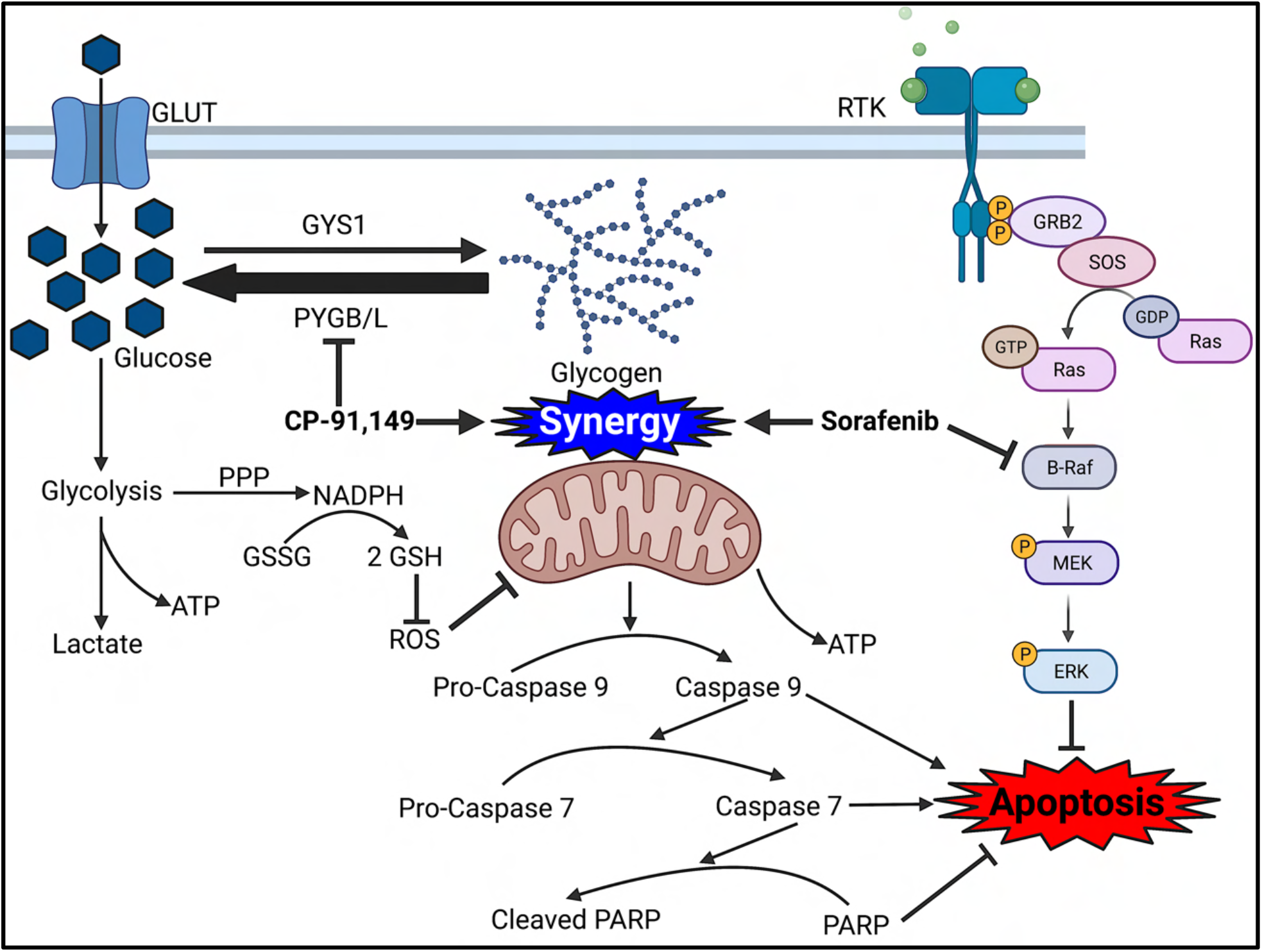
Glycogen phosphorylase inhibition inhibits the pentose phosphate pathway in ATC cells to induce ROS accumulation, mitochondrial dysfunction, and apoptosis. CP-91,149 treatment causes a buildup of glycogen and the glycogen intermediate glucose-1-phosphate. Glucose transporter expression is increased, causing increased glucose flux into the cell to power glycolysis and lactate production and export. The pentose phosphate pathway is impaired to glycogenolysis inhibition, causing a decrease in levels of NADPH and reduced glutathione (GSH). This lack of intracellular reducing power causes an increase in reactive oxygen species, a decrease in the rate of oxygen consumption, decreased ATP levels, and induction of apoptosis through caspase 9, caspase 7, and PARP. CP-91,149 displays drug synergy with the BRAF inhibitor sorafenib.

## Methods

Please see the Supplementary Materials and Methods for additional protocols and reagents (Key Resources Table).

### Culture of Thyroid Cell Lines

Unless otherwise stated, cells were cultured in complete thyroid media (CTM): RPMI-1640 growth media with L-glutamine (300 mg/L), sodium pyruvate and nonessential amino acids (1%) (Corning Inc.) supplemented with 10% fetal bovine serum (Peak Serum) and penicillin-streptomycin (200 IU/L) (Corning Inc.) at 37°C, 5% CO_2_, and 100% humidity. All data were generated from cell lines within 1-5 passages from acquisition. See Supplementary Materials and Methods for detailed information on cell line sources and identifiers. All cell lines were tested regularly for *Mycoplasma* via PCR and validated via short-tandem repeat profiling prior to use.

### Immunofluorescent Analysis of Patient-Derived Tissue Microarray

Thyroid tissue microarray slides (US Biomax, Inc, TH641) were baked at 60°C for 90 minutes and deparaffinized with three changes of xylene (Sigma-Aldrich) and two changes each of sequential alcohol dilutions (100%, 95%, 70%, 50%). After a water rinse, slides were immersed in 1X Dako (Agilent Technologies) for 20 minutes at 95°C for antigen retrieval. Cooled slides were washed 3X in PBS and blocked for 1 hour at room temperature in 10% normal goat serum (Jackson ImmunoResearch) diluted in 5% BSA in PBS with 0.3% Triton X-100. Primary antibodies, diluted in PBS containing 10% normal goat serum, 5.0% BSA, and 0.3% Triton X-100, were added and incubated overnight at 4°C. Slides were washed 7X with 5.0% BSA in PBS. Secondary antibodies, diluted in 5.0% BSA in PBS, were applied for 1 hour at room temperature. The TMA slides were washed, stained with 10 μg/mL DAPI (Thermo Fisher Scientific) for 15 minutes, and then washed in 5.0% BSA in PBS. Slides were mounted using Dako mounting medium, and proteins were detected using antibodies optimized for immunofluorescence. Images were captured using a Zeiss LSM510 META confocal microscope and quantified by ImageJ. Slides were then stained using the PAS protocol in the Supplementary Materials and Methods.

### RNA Extraction and Quantitative Real-Time PCR (qRT-PCR)

Total RNA was extracted using RNeasy Plus Kit (Qiagen) according to manufacturer’s protocol. cDNA was synthesized using LunaScript^®^ RT SuperMix Kit (New England Biolabs). Gene expression to validate RNA-seq analysis was quantified by qRT-PCR using Luna^®^ Universal qPCR Master Mix (New England Biolabs) on a QuantStudio 3 real-time PCR system (Thermo Fisher Scientific). Fold change in gene expression compared to endogenous controls was calculated using the ddCT method. Primer sequences (Eurofins Genomics) are indicated in Supplementary Materials and Methods.

### Immunoblot Analysis

Immunoblots were conducted as described previously^61^. Briefly, proteins were isolated from whole cells in lysis buffer (20mM Tris-HCl (pH 8), 137 mM NaCl, 10% glycerol, 1% Triton X-100, and 2mM EDTA), quantified via Pierce™ BCA Protein Assay Kit (Thermo Fisher Scientific), and 25 µg protein/sample were resolved by polyacrylamide gel electrophoresis and immobilized onto nitrocellulose membranes. Membranes were blocked with 5% w/v BSA in TBS and 0.1% v/v Tween20 (Gold Bio, St Louis, MO, USA) for one hour at room temperature and incubated with primary antibodies overnight. Membranes were washed with TBS-0.1% Tween20 and secondary antibodies were applied for 1 hour at room temperature in 5% w/v BSA in TBS and 0.1% v/v Tween20. Immunoreactive proteins were detected by enhanced chemiluminescence (Thermo Scientific) on a ChemiDoc XRS+ (Bio-Rad Laboratories). Densitometry analysis was performed using ImageJ.

### Quantification of Levels of Glycogen, Glucose-1-Phosphate, Glucose-6-Phosphate, Lactate, ATP, Glucose, NADPH, and Glutathione

Briefly, cells were seeded in 10 cm dishes (for measuring glycogen and NADPH), 6 well plates (G1P, G6P, lactate, glucose, and glutathione), or 96 well plates (ATP) and allowed to adhere overnight. CTM was aspirated and cells were rinsed with PBS before application of compounds dissolved in CTM for 24 hours, at which point cells were ∼90% confluent. CTM was then aspirated, cells were rinsed with PBS, and lysed according to the respective manufacturer protocol (see Key Resources table) for metabolite analysis.

### *In vitro* Cell Staining for Measuring Drug Efficacy

Thyroid cells (4.0 × 10^3^) were seeded in 96 well plates with 100 µl CTM and allowed to adhere overnight. 100 µl of CTM containing 2X concentration of compound was overlayed in each well for 48 hours unless otherwise indicated. Each well then received 100 μl of 10% w/v cold trichloroacetic acid (Fisher) and incubated at 4°C for 1 hour to fix the cells. Each well was rinsed 3X with DI H_2_O and stained with 100 μl 0.057% w/v sulforhodamine B (SRB, Sigma-Aldrich) for 30 minutes at room temperature in the dark. Wells were washed 4X with 300 ul 1% v/v acetic acid (Fisher), and remaining SRB stain was solubilized with 200 μl of 10 mM Tris buffer, pH 10.5 (Fisher). Absorbance was measured at 564 nm with a Synergy 2 Multi-Detection Microplate Reader (Agilent Technologies) and normalized to a vehicle control. Drug synergy was evaluated using the coefficient of drug interaction (CDI) formula: CDI = AB / (A x B) X 100, where AB is the percent remaining cells of an indicated combination treatment, A is the average percent remaining cells of agent 1 alone, and B is the average percent remaining cells of agent 2 alone. A CDI of ≤ 0.7 is considered significantly synergistic; CDI = 1 is additive; CDI > 1.0 is antagonistic^62^.

### *PYGB* RNA Interference

For validating successful *PYGB* knockdown, 4.0 × 10^4^ cells were plated in 12 well plates containing 0.2% Lipofectamine™ 3000 Transfection Reagent and the indicated concentration of siRNA targeting *PYGB* or control siRNA in 1 ml CTM. RNA was extracted 72 hours later for RT-qPCR analysis and protein was extracted 96 hours later for immunoblotting. For measuring the effect of *PYGB* knockdown on cell viability, 1.0 × 10^3^ cells were seeded in 96 well plates with 0.15% Lipofectamine™ 3000 Transfection Reagent and the indicated concentration of siRNA targeting *PYGB* or control siRNA in 200 μl CTM for 7 days before conducting the SRB assay. For measuring the effect of *PYGB* knockdown on glycogen content, 6.5 × 10^5^ cells were seeded in 10 cm dishes with 0.2% Lipofectamine™ 3000 Transfection Reagent and the indicated concentration of siRNA targeting *PYGB* or control siRNA in 10 ml CTM for 4 days preceding cell lysis for glycogen determination.

### Metabolic Flux Analysis

Thyroid cells (5.0 × 10^4^) were plated in a 96 well Seahorse culture plate with compounds of interest diluted in 200 μl CTM for 24 hours. Cells were washed with PBS and equilibrated for 1 hour at 37°C in a non-CO_2_ incubator with compounds of interest diluted in Seahorse RPMI (supplemented with 11.1 mM glucose, 2 mM glutamine, and 5% serum) before analysis of OCR and ECAR in a Metabolic Flux Analyzer (Seahorse Bioscience, North Billerica, MA 96XP).

### Quantification of Reactive Oxygen Species

Thyroid cells (1.0-2.0 × 10^4^) were seeded in a black 96 well plate in 200 μl CTM and allowed to adhere overnight. Wells were washed with PBS and incubated with or without 25 μM H_2_DCFDA for 45 minutes at 37°C in a non-CO_2_ incubator with compounds of interest diluted in phenol-red free RPMI (supplemented with 2 mM glutamine and 10% charcoal-stripped serum). Wells were washed with PBS and received compounds of interest diluted in phenol-red free, supplemented RPMI and incubated at 37°C in a SpectraMax M4 Microplate Reader (Molecular Devices) to record fluorescent measurements (Ex/Em = 485/535 nm) every 15 minutes overnight. Maximum fluorescence intensity was determined (∼6 hours), background signal (-H_2_DCFDA) was subtracted from each group, and corrected values were normalized to the vehicle control.

### *In vivo* Evaluation of CP-91,149 and Sorafenib

The xenograft experiment was approved by the Animal Care and Use Committee of the University of Vermont (Protocol X0-018). Four-week-old athymic female nude mice (outbred homozygous nude *Foxn1*^*nu*^*/Foxn1*^*nu*^) were purchased from Jackson Laboratory (Bar Harbor, ME, USA) and were allowed to acclimatize for one week. Mice were given food and water *ad libitum*. Mice were anesthetized using 3% isoflurane delivered at 1.5 L/min, and tumors were established by subcutaneously injecting 1 × 10^6^ 8505C cells with a 26G needle in 100 μL of high concentration Matrigel (Corning Inc.) diluted 1:2 with base RPMI-1640 into each flank of 24 mice. One week later, mice were sorted into four treatment groups to achieve approximately equal body weights between mice. Mice were then administered 50 mg/kg CP-91,149, 10 mg/kg sorafenib, both CP-91,149 and sorafenib, or vehicle control every day for 6-7 days a week via intraperitoneal injection using 27G needles. These doses were based on previous reports^32,63,64^. Pharmacological agents were dissolved in 100% DMSO and diluted daily in 30% DMSO, 40% PEG300, and 30% PBS and incubated at 55°C for 10 minutes prior to vortexing to encourage complete solubilization. Tumor dimensions were measured with digital calipers, and the volumes were calculated by the following formula: (**Π** x a x b^2^) / 6, where a represents the largest diameter and b is the perpendicular diameter. The body weight of each animal was taken twice a week to measure toxicity. Mice were euthanized with carbon dioxide, and the tumors were harvested, fixed with formalin (Thermo Fisher Scientific), and stored at 4°C prior to slide sectioning and immunofluorescence analysis.

### Statistical Analysis

All statistical analyses were performed using GraphPad Prism software. Paired comparisons were conducted by Student’s t-test. Group comparisons were made by one-way ANOVA followed by Dunnett’s or Tukey’s multiple comparison test as appropriate. Two-way ANOVA followed by a Tukey’s multiple comparison test was conducted for multigroup analysis. Data are represented as mean ± standard deviation. Area under the curve (AUC) at the 95th confidence interval was used to evaluate statistical differences in indicated assays.

## Data Availability

The data generated in this study are available within the article and its supplementary data files.

## Authors’ Contributions

**C. D. Davidson**: Conceptualization, investigation, writing–original draft, review, and editing, data curation, formal analysis. **J.A. Tomczak**: Writing–review and editing, data curation. **E. Amiel**: Conceptualization, writing–review and editing, resources, funding acquisition, methodology. **F.E. Carr**: Data curation, writing–review and editing, resources, funding acquisition, methodology.

## Acknowledgments

Imaging work was performed at the Microscopy Imaging Center at the University of Vermont. Confocal microscopy was performed on a Zeiss 510 META laser scanning confocal microscope supported by NIH Award Number 1S10RR019246 from the National Center for Research Resources. We are especially grateful for the work performed by Nicole Bouffard and Michele von Turkovich in confocal microscopy and TEM, respectively. TPC-1 cells were generously provided by Dr. John A. Copland III, Mayo Clinic, Jacksonville, FL. B-CPAP, FTC-133, CUTC61, OCUT-2, and 8505C cells were purchased from the CU Denver Cell Technologies Shared Resource’s Cell Line Repository. Human cell line authentication and molecular imaging were performed in the Vermont Integrative Genomics Resource supported by the University of Vermont (UVM) Cancer Center, and the UVM Larner College of Medicine. Additional human cell line authentication was performed by the CU Cancer Center Tissue Culture Shared Resource supported by the National Cancer Institute (P30CA046934). We thank the Histology Research Support Facility for histological services in the Department of Pathology and Laboratory Medicine at the University of Vermont Medical Center, Burlington, VT. We thank the Office of Animal Care Management for their expertise for *in vivo* experiments. We thank Dr. Jane Lian, Professor of Biochemistry, University of Vermont, for her invaluable comments and discussion in preparation for the *in vivo* study.

