## Supplemental Materials for "Inhibition of Glycogen Metabolism Induces Reactive Oxygen Species-Dependent Apoptosis in Anaplastic Thyroid Cancer"

### Supplementary Materials and Methods

#### Periodic Acid Schiff Staining on Patient-Derived Tissue Microarray

After immunofluorescent imaging, TMA slides were rinsed with $H_2O$ and stained for glycogen using the Periodic Acid Schiff (PAS) Stain Pack (Tyr Scientific).

#### Transmission Electron Microscopy of Glycogen Deposits

Cells were seeded in 10 cm dishes and allowed to adhere overnight. CTM was aspirated and cells were rinsed with PBS before application of RPMI with no glucose, 11.1 mM glucose, or 11.1 mM glucose + 50 µM CP-91,149 for 24 hours. Cells were then rinsed with PBS and lifted with trypsin for preparation for TEM as described previously[1].

#### In Vitro Determination of Cell Attachment Potential

8505C cells were treated with or without 50 µM CP-91,149 for 24 hours in T75 flasks. Cells were washed with PBS and lifted with trypsin. $2.5 \times 10^4$ cells were seeded in 200 µl of CTM with or without 50 µM CP-91,149 in 96 well plates for the indicated time points preceding staining with SRB to determine relative number of cells able to attach in a short time frame.

#### Cell Viability Assay for Determination of Live/Dead Cell Ratios

$1.5 \times 10^4$ cells were plated in 12 well tissue culture dishes. After adhering overnight, the cells were treated with an inhibitor or vehicle at the indicated concentrations. Every day after treatment for four days, the media were removed, cells were washed with PBS, lifted with trypsin, and diluted 1:4 into trypan blue. The number of surviving cells was counted with a hemocytometer.

#### Determination of CP-91,149 Efficacy on ATC Stem Cells

ATC thyrospheres were grown and counted as described previously using RPMI-1640 supplemented with 20 ng/mL each of epidermal growth factor (EGF) and fibroblastic growth factor (FGF2) (GoldBio)[2]. Thyrospheres per well were counted and imaged using a digital camera (Diagnostics Instruments) connected to a Nikon Eclipse TS100 inverted microscope.

**Key Resources Table**

| Reagent or Resource | Source | Identifier |
|---|---|---|
| **Antibodies (application & dilution)** | | |
| Anti-GYS1 (WB, 1:10000 and IF, 1:500) | Abcam | Cat# ab40810; RRID:AB_732660 |
| Anti-pGYS1 (WB, 1:1000) | Millipore Sigma | Cat# 07-817; RRID:AB_568824 |
| Anti-PYGL (WB, 1:1000 and IF, 1:500) | Abcam | Cat# ab198268; RRID:AB_2894845 |
| Anti-PYGB (WB, 1:1500 and IF, 1:500) | Invitrogen | Cat# PA5-28022; RRID:AB_2545498 |
| Pro and cleaved PARP (WB, 1:1000) | Cell Signaling Technology | Cat# 9542; RRID:AB_2160739 |
| Pro and cleaved Caspase 7 (WB, 1:500) | Cell Signaling Technology | Cat# 9494; RRID:AB_2068141 |
| Pro and cleaved Caspase 9 (WB, 1:1000) | Cell Signaling Technology | Cat# 9502; RRID:AB_2068621 |
| Ki-67 (WB, 1:10000 and IF, 1:500) | Thermo Fisher Scientific | Cat# PA5-116479; RRID:AB_2894846 |
| GAPDH (WB, 1:10000) | Rockland Immunochemicals | Cat# 600-401-A33; RRID:AB_2107593 |
| β-actin (WB, 1:3000) | Thermo Fisher Scientific | Cat# MA5-15739; RRID:AB_10979409 |
| Cleaved PARP (IF 1:100) | Cell Signaling Technology | Cat# 9544; RRID:AB_2160724 |
| Horse Anti-Mouse IgG HRP Conjugate (WB, 1:10000) | Cell Signaling Technology | Cat# 7076; RRID:AB_330924 |
| Goat Anti-Rabbit IgG HRP Conjugate (WB, 1:10000) | Cell Signaling Technology | Cat# 7074; RRID:AB_2099233 |

| | | |
|---|---|---|
| Goat Anti-Rabbit IgG Alexa Fluor® 488 Conjugate (IF, 1:500) | Cell Signaling Technology | Cat# 4412; RRID:AB_1904025 |
| Goat Anti-Mouse IgG Alexa Fluor® 594 Conjugate (IF, 1:500) | Cell Signaling Technology | Cat# 8890; RRID:AB_2714182 |
| **Media, Supplements, Small Molecule Reagents, and Cell Culture Supplies** | | |
| RPMI (glucose and glutamine replete) | Sigma-Aldrich | Cat# R8758 |
| Sodium pyruvate | Corning Inc. | Cat# 25-000-CI |
| Nonessential amino acids | Corning Inc. | Cat# 25-025-CI |
| Fetal bovine serum | Peak Serum | Cat# PS-FB2 |
| Charcoal stripped serum | Sigma-Aldrich | Cat# F6765 |
| Penicillin-Streptomycin | Corning Inc. | Cat# 30-002-CI |
| RPMI (No glucose, No pyruvate, glutamine replete) | Gibco | Cat# 11879020 |
| RPMI (No glucose, No pyruvate, No glutamine, No phenol red) | Gibco | Cat# A2494201 |
| Agilent Seahorse XF RPMI | Agilent Technologies | Cat# 103576-100 |
| Glutamine | Sigma-Aldrich | Cat# G7513 |
| Glucose | Research Products International | Cat# G32040 |
| Recombinant Human Fibroblast Growth Factor 2 | GoldBio | Cat# 1140-02-10 |
| Recombinant Human Epidermal Growth Factor | GoldBio | Cat# 1150-04-100 |
| DMSO | Sigma-Aldrich | Cat# D2650 |
| Trypsin-EDTA | Gibco | Cat# 25200056 |
| PBS | Corning, Inc. | Cat# 21-040-CV |
| Trypan blue | Corning, Inc. | Cat# 25-900-CI |
| Buparlisib | MedChemExpres s | Cat# Hy70063 |
| Alpelisib | MedChemExpres s | Cat# Hy-15244 |

| | | |
|---|---|---|
| Sorafenib | MedChemExpres s | Cat# Hy-10201 |
| Lapatinib | MedChemExpres s | Cat# Hy-50898 |
| Palbociclib | MedChemExpres s | Cat# Hy50767 |
| Lenvatinib | MedChemExpres s | Cat# HY-10981 |
| Doxorubicin | Abcam | Cat# ab120629 |
| CP-91,149 | Selleckchem | Cat# S2717 |
| CP 316819 | Tocris Bioscience | Cat# 3542 |
| 6-AN | Cayman Chem | Cat# 10009315 |
| 2-DG | Selleckchem | Cat# S4701 |
| 3-BP | Cayman Chem | Cat# 19068 |
| Guaiacol | Acros Organics | Cat# AC120192500 |
| yGsy2p-IN-1 | MedChemExpres s | Cat# HY-131062 |
| MnTMPyP | Sigma-Aldrich | Cat# 475872 |
| N-Acetylcysteine | MedChemExpres s | Cat# HY-B0215 |
| $H_2$DCFDA | Selleckchem | Cat# S9687 |
| NADPH | Santa Cruz Biotechnology | Cat# SC-202725 |
| 30% $H_2O_2$ | Sigma-Aldrich | Cat# 216763 |
| High concentration basement membrane matrix Matrigel® | Corning, Inc. | Cat# 354248 |
| Polyethylene glycol (PEG) 300 | Sigma-Aldrich | Cat# 202371 |
| **Critical Commercial Assays** | | |
| RNeasy Mini Kit (RNA extraction) | Qiagen | Cat# 74106 |
| LunaScript® RT SuperMix Kit | New England Biolabs | Cat# E3010 |
| Luna® Universal qPCR Master Mix | New England Biolabs | Cat# M3003 |
| Lipofectamine™ 3000 Transfection Reagent | Thermo Fisher Scientific | Cat# L3000015 |
| Silencer Select Negative Control siRNA | Invitrogen | Cat# 4390844 |

| | | |
|---|---|---|
| Silencer ® Select siRNA for *PYGB* | Thermo Fisher Scientific | Assay ID: s11641 |
| Glycogen Colorimetric/Fluorometric Assay Kit | BioVision Inc. | Cat# K646-100 |
| Pierce™ BCA Protein Assay Kit | Thermo Fisher Scientific | Cat# 23225 |
| Glucose-1-Phosphate Colorimetric Assay Kit | BioVision Inc. | Cat# K697 |
| EnzyChrom™ GSH/GSSG Assay Kit | BioAssay Systems | Cat# EGTT-100 |
| EnzyChrom™ L-Lactate Assay Kit | BioAssay Systems | Cat# ECLC-100 |
| Glucose Assay Kit I | Eton Bioscience | Cat# 1200031002 |
| CellTiter-Glo® 2.0 Cell Viability ATP Assay | Promega | Cat# G9241 |
| NADPH Assay Kit | Abcam | Cat# ab186031 |
| **Staining Reagents** | | |
| Periodic Acid Schiff (PAS) Stain Pack | Tyr Scientific | Cat# TS9-500PK |
| Trichloroacetic acid | Fisher Bioreagents | Cat# BP555-250 |
| Sulforhodamine B | Sigma-Aldrich | Cat# S1402 |
| Glacial acetic acid | Fisher Bioreagents | Cat# BP2401-500 |
| Tris base | Fisher Bioreagents | Cat# BP152-1 |
| Formal-Fixx™ formalin | Thermo Fisher Scientific | Cat# 9990244 |
| Xylene | Sigma-Aldrich | Cat# 214736 |
| Ethyl alcohol, 200 proof | Pharmco | Cat# 111000200 |
| Dako Antigen Retrieval, pH 6.1 | Agilent Technologies | Cat# S169984-2 |
| Normal goat serum | Jackson ImmunoResearch | Cat# 005-000-121, RRID: AB_2336990 |
| DAPI | Thermo Fisher Scientific | Cat# D1306 |
| Dako Fluorescence Mounting Medium | Agilent Technologies | Cat# S3023 |
| **Cell Lines** | | |
| Nthy-ori 3-1 | Sigma-Aldrich | Cat# 90011609, RRID:CVCL_2659 |

| | | |
|---|---|---|
| TPC-1 | Mayo Clinic, Jacksonville FL, USA | Marlow, et al., *J Clin Endocrinol Metab*, 2018. |
| B-CPAP, FTC-133, CUTC61, OCUT-2, 8505C | University of Colorado, Denver CO, USA | Landa, et al., *Clin Cancer Res*, 2019. |
| **Mouse Model** | | |
| 4 wk female J:NU *Foxn1*$^{nu}$/*Foxn1*$^{nu}$ | Jackson Laboratory | Cat# 007850, RRID:IMSR_JAX:007850 |
| **Software** | | |
| ImageJ | Open Source/National Institutes of Health | https://imagej.nih.gov/ij/ |
| GraphPad Prism | GraphPad Software, Inc. | https://www.graphpad.com/scientific- software/prism/ |
| Image Studio™ | LI-COR | https://www.licor.com/bio/image-studio-lite/download |
| NIS-Elements | Nikon Instruments | https://www.microscope.healthcare.nikon.com/products/software/nis-elements |
| Gen5 Data Analysis | BioTek | https://www.biotek.com/products/software-robotics-software/gen5-microplate-reader-and-imager-software/ |
| Seahorse Wave Software | Agilent Technologies | https://www.agilent.com/en/product/cell-analysis/real-time-cell-metabolic-analysis/xf-software/seahorse-wave-desktop-software-740897 |

**Oligonucleotides**

| **Target** | **Forward (5'-3')** | **Reverse (5'-3')** | **Amplicon size (bp)** |
|---|---|---|---|
| *GYS1* | ACAACCTGGAGAACTTCAAC | ATCTGGGACACAGTAGTGAA | 118 |
| *PYGL* | TGCCCGGCTACATGAATAACA | TGTCATTGGGATAGAGGACCC | 162 |
| *PYGB* | ACGCAGCAGCACTACTAC | TCGCAGGCATTCTGAAGG | 119 |
| *SLC2A1* | TGGCATCAACGCTGTCTTCT | AGCCAATGGTGGCATACACA | 83 |
| *SLC2A3* | CTGAGGACGTGGAGAAAACTTG | AATATCAGAGCTGGGGTGACCTTC | 154 |
| *SLC2A4* | CAGTGGCTTGGAAGGAAAAGG | CAGGTGAGTGGGAGCAATCT | 195 |

**Supplementary Table S1.** Genetic Background of Thyroid Cell Lines[3]

| | **Cell Line:** | **Nthy-ori-3-1** | **TPC-1** | **B-CPAP** | **FTC-133** | **CUTC61** | **OCUT-2** | **8505C** |
|---|---|---|---|---|---|---|---|---|
| | **Tumor Type:** | Normal | PTC | PTC | FTC | FTC | ATC | ATC |
| **Gene Function** | **Gene Name** | **Mutation Type** | | | | | | |
| MAPK Signaling | *BRAF* | WT | WT | **Missense** | WT | WT | **Missense** | **Missense** |
| Telomerase | *TERT* (promoter) | WT | 124C-T | 124C-T 125C-T | 124C-T | 124C-T | 146C-T | 146C-T |
| DNA Homeostasis | *TP53* | WT | WT | Missense | **Missense** | Truncated | WT | Missense |
| PI3K Signaling | *PIK3CA* | WT | WT | WT | WT | WT | Missense | WT |
| PI3K Signaling | *PTEN* | WT | WT | WT | **Truncated** | WT | WT | WT |
| PI3K and MAPK Signaling | *CCDC6-RET* | WT | **Fusion** | WT | WT | WT | WT | WT |
| PI3K and MAPK Signaling | *NF1* | WT | WT | WT | **Truncated** | WT | WT | WT |
| PI3K and MAPK Signaling | *HRAS* | WT | WT | WT | WT | **Missense** | WT | WT |
| Cell Cycle | *P16* | WT | Truncated | WT | WT | WT | WT | WT |
| Cell Cycle | *RB1* | WT | WT | WT | In Frame | WT | WT | WT |
| DNA Homeostasis | *ATM* | WT | WT | WT | WT | WT | WT | Truncated |
| DNA Mismatch Repair | *MLH1* | WT | WT | Missense | WT | WT | WT | WT |
| DNA Mismatch Repair | *MSH6* | WT | WT | WT | Truncated | WT | WT | WT |

**Bold** = Driver Mutation(s)

### Supplementary Figures

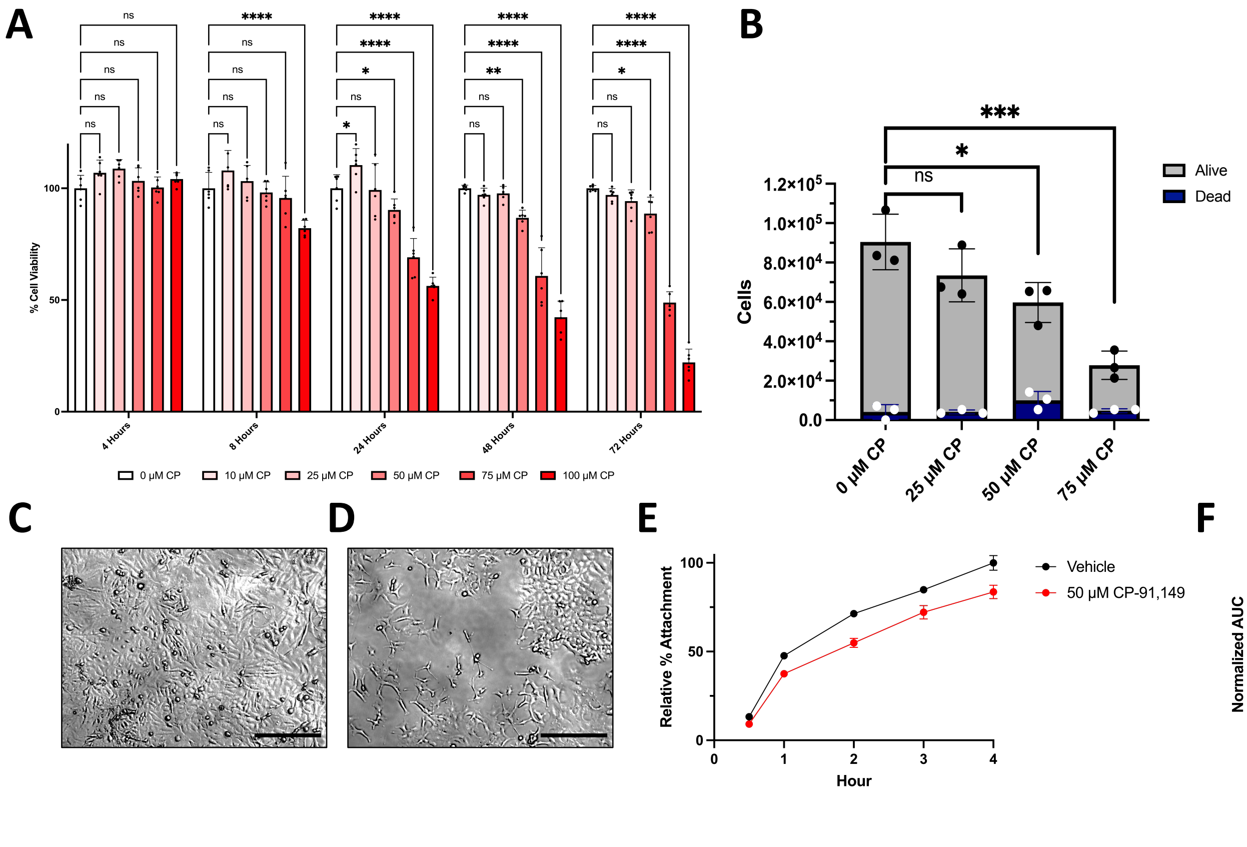


**Supplementary Figure S1.** CP-91,149 decreases 8505C cell viability in a time-dependent manner and inhibits cell attachment.

**A.** 8505C cells were treated with CP-91,149 at various concentrations and time points for SRB assay. **B.** 8505C were stained with trypan blue and hand counted for dead and alive cells after 48 hours of CP-91,149 incubation at the indicated concentrations. **C-D.** Cells were imaged 48 hours following vehicle (C) or CP treatment (D). Magnification = 40X, scalebar = 100 µM **E.** Cells were challenged to adhere to fibronectin-coated cell culture plates following 24 hours of 50 uM CP treatment, as measured by SRB assay. **F.** Area under the curve analysis of cell attachment assay.

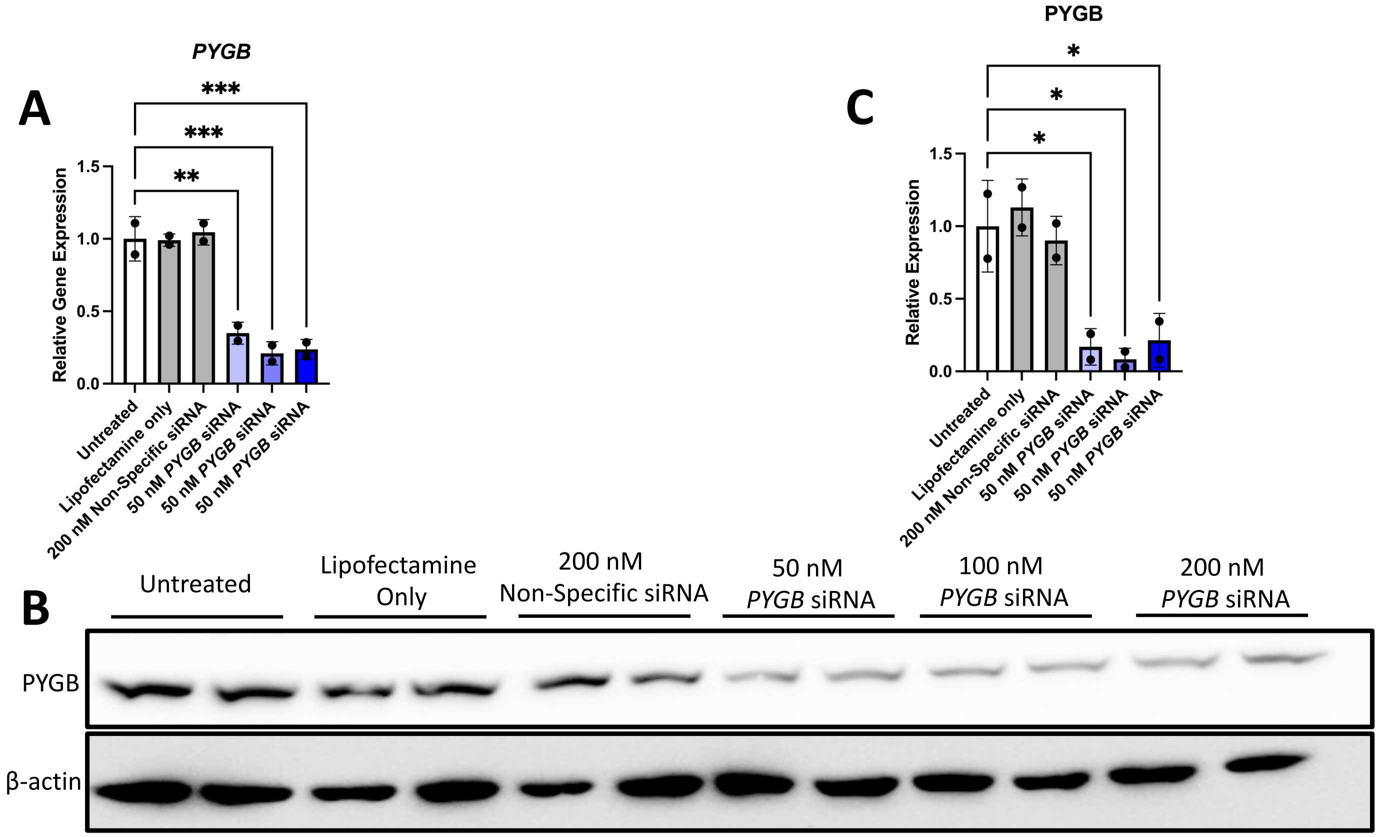


**Supplementary Figure S2.** Validation of *PYGB* knockdown.

**A-B.** 8505C cells were reverse transfected using lipofectamine with antisense *PYGB* siRNA at the indicated concentrations for 72 hours for RT-qPCR analysis of *PYGB* expression (A) and 96 hours for immunoblot analysis of PYGB protein (B). **C.** Densitometry analysis of PYGB immunoblot.

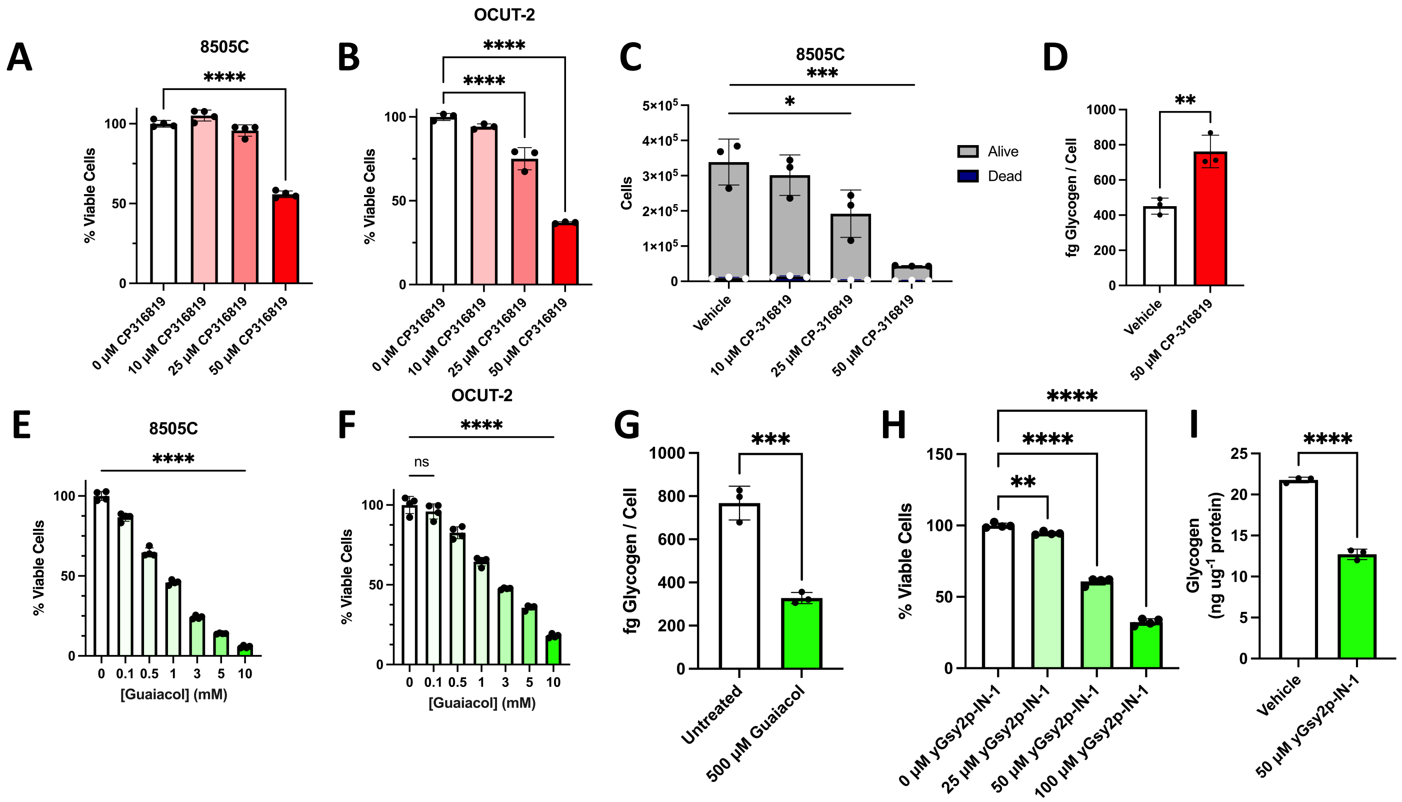


**Supplementary Figure S3.** Glycogen metabolism is inhibited with various small molecule inhibitors in ATC cells.

**A-C.** Cell viability was determined via SRB assay (A-B) and hand counting with trypan blue (C) following treatment with CP316819 at the indicated concentrations for 48 hours. **D.** Glycogen assay was conducted in 8505C cells following overnight treatment with CP-316819. **E-F.** Cell viability was determined via SRB assay in 8505C (E) and OCUT-2 (F) cells following treatment with guaiacol at the indicated concentrations for 48 hours. **G.** Glycogen assay was conducted in 8505C cells following overnight treatment with guaiacol. **H.** Cell viability was determined via SRB assay in 8505C following treatment with yGsy2p-IN-1 at the indicated concentrations for 48 hours. **I.** Glycogen assay was conducted in 8505C cells following overnight treatment with yGsy2p-IN-1.

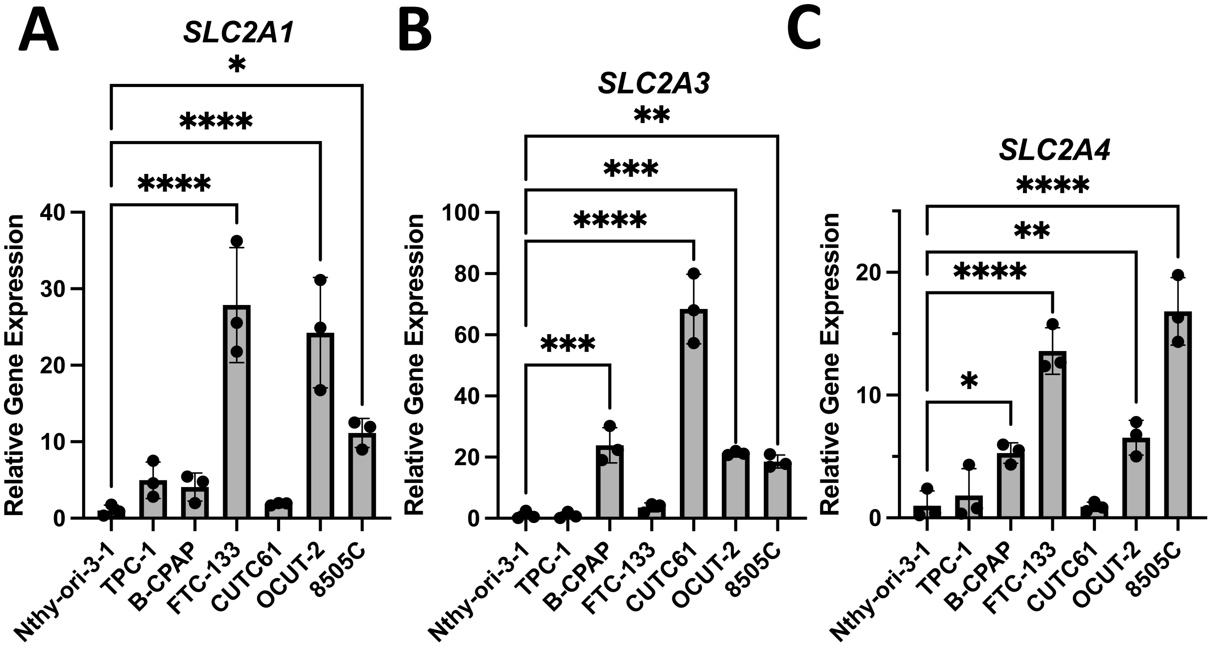


**Supplementary Figure S4.** Baseline transcript levels of glucose transporters in thyroid cancer cell lines.

**A-C.** Thyroid cells were cultured in normal media conditions and analyzed for expression of *SLC2A1* (A), *SLC2A3* (B), and *SLC2A4* (C) using RT-qPCR.

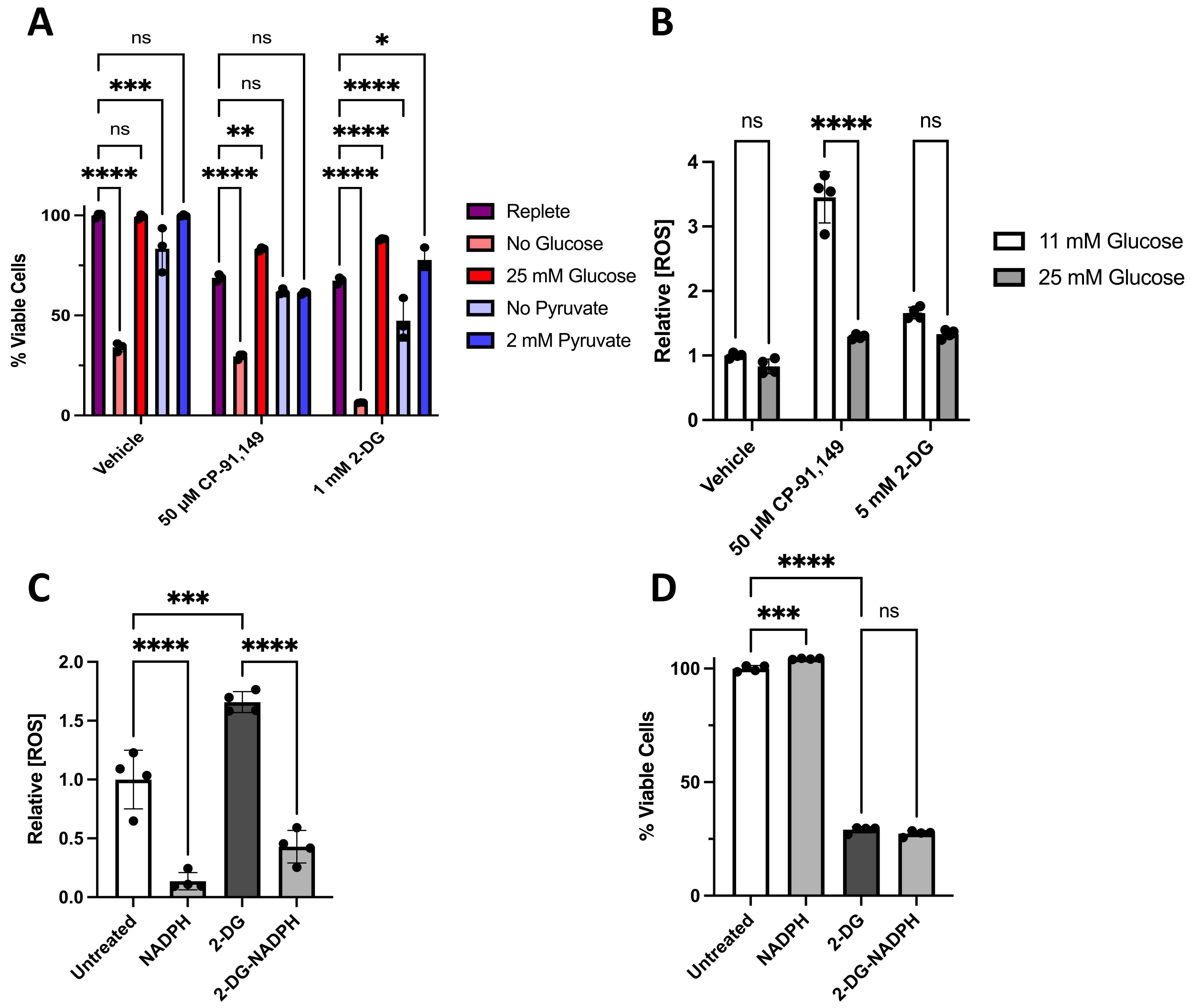


**Supplementary Figure S5.** Glucose and pyruvate availably selectively modulate ATC cell susceptibility to glycolytic and glycogenolytic inhibition.

**A.** 8505C cells were treated with CP or 2-DG for 48 hours in the indicated cell culture media, and cell viability was determined via SRB assay. **B.** 8505C cells were treated with CP or 2-DG in the indicated cell culture media prior to ROS determination. **C.** 8505C cells were treated with NADPH, 2-DG, or both immediately prior to ROS assay in replete media. **D.** 8505C cells were treated with NADPH, 2-DG, or both for 48 hours in replete media prior to SRB assay.

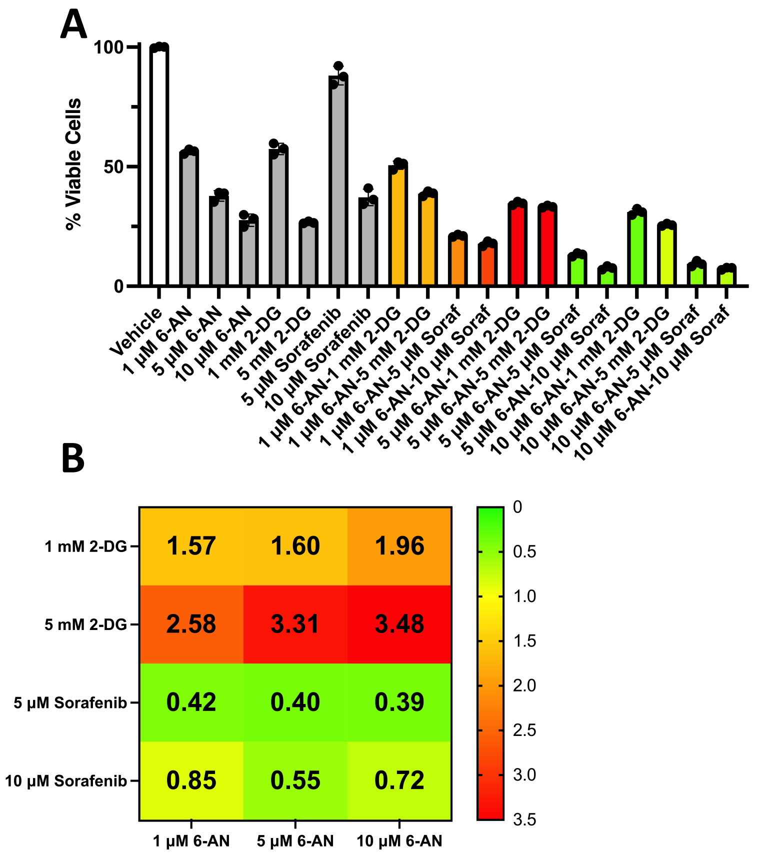


**Supplementary Figure S6.** Sorafenib is synergistic with inhibition of the PPP but not glycolysis. **A-B.** SRB assay was conducted on 8505C cells treated with the indicated combinations of inhibitors for 48 hours (A) to calculate CDI values (B).
